# Repurposing a biosynthetic enzyme for imaging UDP-glucose dynamics in living systems

**DOI:** 10.64898/2026.11.13.766517

**Authors:** Zefan Li, Delai Huang, Oluwaseun Mustapha, Franklin Herbert, Adam Thuen, Sijie Hao, David M. Parichy, Mikhail Drobizhev, Hui-wang Ai

## Abstract

Metabolite dynamics remain difficult to visualize in living systems, particularly for nucleotide sugars that link metabolism to glycosylation and signaling. Here we repurpose *Helicobacter pylori* UDP-glucose pyrophosphorylase to develop UGluS, a genetically encoded ratiometric fluorescent sensor for its biosynthetic product, UDP-glucose (UDP-Glc). Circularly permuted GFP insertion and directed evolution yield an approximately 18-fold excitation-ratio response and an apparent dissociation constant of 32.5 μM, with preferential responsiveness to UDP-Glc over related metabolites. Live-cell imaging and microplate assays reveal responses to metabolic, genetic and mechanosensitive perturbations in mammalian cells. In zebrafish larvae, UGluS reveals uridine-induced increases in UDP-Glc availability and fin-fold enrichment. In mouse liver slices and skeletal muscle in vivo, insulin- and glucose-induced decreases in UDP-Glc availability accompany increased glycogen content. These findings establish UGluS as a tool for imaging UDP-Glc dynamics across biological scales and extend the use of biosynthetic enzymes as fluorescent sensing scaffolds.

---

Cellular function depends on the coordinated production, utilization, and distribution of metabolites. Although chromatographic and mass spectrometric methods enable metabolite quantification, conventional bulk analyses rely on destructive sampling and extraction, providing discrete snapshots that obscure spatial heterogeneity and constrain the temporal resolution of dynamic measurements.^1,2^ Genetically encoded fluorescent biosensors complement these approaches by enabling spatially and temporally resolved monitoring of metabolites in living cells and organisms, but currently cover only a small fraction of cellular metabolites.^3^ Expanding this repertoire requires identifying suitable molecular-recognition scaffolds and coupling metabolite binding to a robust fluorescence response—two interrelated challenges that remain major barriers to sensor development.

Enzymes offer a diverse pool of molecular-recognition scaffolds, and previous studies have successfully repurposed enzyme-derived recognition domains to develop fluorescent metabolite sensors.^4–7^ Although these sensors have employed various substrate- and cofactor-recognition mechanisms, harnessing the recognition of a biosynthetic product by the enzyme responsible for its synthesis remains a relatively underexplored strategy. This approach requires preserving product recognition while suppressing catalytic activity, minimizing interference from substrates, cofactors and related metabolites, and coupling ligand binding to a robust fluorescence response. Applying this strategy to a nucleotide-sugar biosynthetic enzyme could expand the repertoire of genetically encoded metabolite sensors while highlighting the broader potential of product-binding enzymes as sensing scaffolds.

The nucleotide sugar UDP-glucose (UDP-Glc) offers an important target for this approach. Synthesized from glucose-1-phosphate and UTP by UDP-glucose pyrophosphorylase (UGP), UDP-Glc connects carbohydrate metabolism and nucleotide availability to downstream biosynthesis and signaling (**Fig. 1A**).^8^ It supplies glucose for glycogen synthesis and contributes to glycan and extracellular matrix biosynthesis, thereby linking nutrient availability to carbon storage and cellular architecture.^9–12^ UDP-Glc has also been implicated in the regulation of lipogenesis, protein quality control, and extracellular signaling (**Fig. 1B**).^13–21^ Disruptions of UDP-Glc metabolism have been associated with neurological disease, cancer, and kidney injury.^1,22–27^ Despite these diverse functions, the spatial and temporal behavior of UDP-Glc remains difficult to interrogate directly in intact biological systems. This limits our understanding of how rapidly UDP-Glc pools respond to changing metabolic and physiological inputs and how their availability varies across tissues.

**Figure 1.**
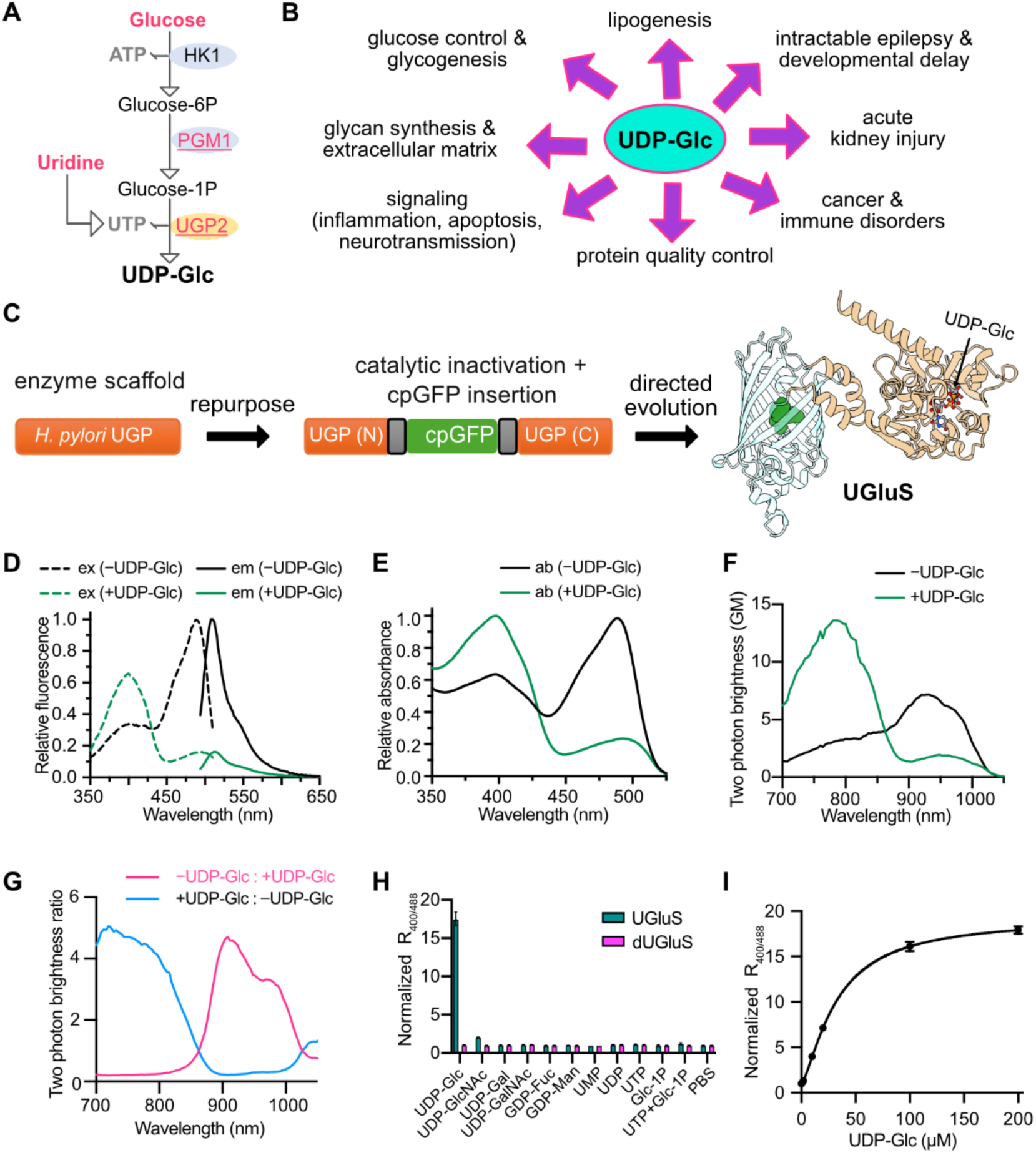
UDP-glucose biology and development and *in vitro* characterization of the enzyme-derived sensor UGluS. **(A)** Schematic of UDP-glucose (UDP-Glc) biosynthesis. HK1, hexokinase 1; PGM1, phosphoglucomutase 1; UGP2, UDP-glucose pyrophosphorylase 2. **(B)** Overview of the biological functions and disease associations of UDP-Glc. **(C)** Engineering strategy for UGluS. A *Helicobacter pylori* UDP-glucose pyrophosphorylase (UGP) scaffold was catalytically inactivated, modified by insertion of a circularly permuted GFP (cpGFP), and optimized through directed evolution. UGP (N) and UGP (C) denote the N- and C-terminal segments of the UGP scaffold, respectively. The resulting sensor architecture is illustrated on the right. **(D)** Fluorescence excitation (ex; dashed lines) and emission (em; solid lines) spectra of purified UGluS in the absence (black) and presence (green) of 100 μM UDP-Glc. **(E)** Absorbance spectra of purified UGluS in the absence (black) and presence (green) of 100 μM UDP-Glc. **(F)** Two-photon excitation brightness spectra of purified UGluS in the absence (black) and presence (green) of 100 μM UDP-Glc. **(G)** Wavelength-dependent two-photon brightness ratios calculated from the spectra in (**F**). Magenta and blue curves show the ratios −UDP-Glc/+UDP-Glc and +UDP-Glc/−UDP-Glc, respectively, at each excitation wavelength. **(H)** Selectivity profiles of UGluS and the unresponsive control variant dUGluS for UDP-Glc and related metabolites, measured as normalized fluorescence excitation ratios (400/488 nm). PBS, phosphate-buffered saline. **(I)** UDP-Glc concentration–response curve of UGluS, measured as normalized fluorescence excitation ratios (400/488 nm). The data were fitted with the Hill equation to estimate the apparent dissociation constant (*K*_d_ = 32.5 ± 1.4 μM) and Hill coefficient (*h* = 1.4 ± 0.1). Fitted parameters are reported as best-fit estimates ± standard error. Data in **D–G** are representative spectra and derived spectral ratios from three independent experiments that yielded similar results. Data in **H** and **I** are presented as mean ± SD (*n* = 3 technical replicates). Ratios in H and I were normalized to the corresponding ligand-free values.

A genetically encoded fluorescent sensor would enable UDP-Glc dynamics to be followed in living cells, tissues, and whole organisms under basal and perturbed conditions. Such a tool would complement bulk measurements by resolving temporal changes and spatial heterogeneity in UDP-Glc availability, providing a means to investigate its regulation in physiological and pathophysiological contexts.

Here we engineer UGluS, a genetically encoded ratiometric fluorescent sensor for UDP-Glc, by inserting circularly permuted green fluorescent protein (cpGFP) into *H. pylori* UGP (HpUGP) and optimizing the resulting construct by directed evolution. In vitro characterization establishes a large, preferential response to UDP-Glc and ligand-dependent fluorescence changes under one- and two-photon excitation. Using UGluS, we resolve UDP-Glc changes in response to metabolic, genetic, and mechanosensitive perturbations in mammalian cells and reveal spatial differences in zebrafish larvae. We further demonstrate imaging of UDP-Glc dynamics in response to physiological stimulation in mouse liver slices and skeletal muscle in live mice. These studies establish a tool for monitoring UDP-Glc dynamics across biological scales and provide a proof of concept for repurposing a nucleotide-sugar biosynthetic enzyme as a fluorescent reporter of its biosynthetic product.

## Results

### Repurposing UDP-glucose pyrophosphorylase as a fluorescent recognition scaffold

To enable direct monitoring of UDP-Glc in living systems, we sought to engineer a genetically encoded fluorescent sensor with high selectivity and a large dynamic range. Our previous sensor for UDP-N-acetylglucosamine (UDP-GlcNAc) used a catalytically inactive glycosyltransferase to recognize its nucleotide-sugar donor substrate but was susceptible to interference from UDP and UTP.^7^ This limitation motivated us to explore a biosynthetic enzyme as an alternative recognition scaffold. We selected UGP, which synthesizes UDP-Glc, to test whether recognition of its biosynthetic product could be coupled to a selective fluorescence response. Specifically, we chose HpUGP because crystal structures of both its apo and UDP-Glc-bound forms are available and the enzyme binds UDP-Glc with micromolar affinity.^28^ To suppress catalysis while retaining UDP-Glc recognition, we introduced the R15A substitution, which was previously reported to suppress HpUGP catalytic activity while preserving UDP-Glc binding.^28^ We then compared the apo and ligand-bound structures to identify regions undergoing ligand-associated conformational rearrangements and selected four candidate sites (residues 68, 80, 135, and 227) for circularly permuted GFP (cpGFP) insertion (**Supplementary Fig. 1A,B**).

For each candidate site, we generated a cpGFP insertion library with one randomized amino acid in each fusion linker (**Supplementary Fig. 1C**). Following insertion-site screening, we advanced the construct with cpGFP inserted between residues 80 and 81 for linker optimization (**Supplementary Fig. 2A,B**). Optimization of linker length and composition using randomized one-, two-, and three-residue linkers yielded UGluS0.1, with an excitation ratio response of approximately 1.5 in the screening assay (**Supplementary Figs. 1C and 2B**). For plate-reader measurements, the response was defined as R/R_0_, where R is the fluorescence excitation ratio at 400 and 488 nm (F_400_/F_488_) in the presence of UDP-Glc and R_0_ is the corresponding ligand-free ratio.

We next subjected UGluS0.1 to iterative directed evolution to improve sensor performance. Sixteen rounds of error-prone PCR and screening produced UGluS0.2, which showed a substantially increased ratiometric response but reduced fluorescence brightness (**Supplementary Figs. 1C and 2C**). Introducing the chromophore-proximal substitutions S139T and S253A, previously used to improve the brightness of cpGFP-based biosensors,^29^ yielded the brighter UGluS0.3 at the expense of its ratiometric response (**Supplementary Figs. 2C and 3**). Three further rounds of directed evolution produced the final sensor, UGluS, which exhibited an approximately 18-fold ligand-dependent change in excitation ratio (**Fig. 1C and Supplementary Table 1**). The complete workflow comprised 19 rounds of error-prone PCR and an intervening brightening step (**Supplementary Fig. 2C**). To generate a ligand-unresponsive control, we introduced the Q350A and T479A substitutions into UGluS to disrupt UDP-Glc binding, yielding dUGluS (**Supplementary Fig. 3**).

### Photophysical characterization and metabolite responsiveness of UGluS

Purified UGluS displayed a pronounced UDP-Glc-dependent change in the relative intensities of its two fluorescence excitation bands (**Fig. 1D**). Photophysical measurements performed independently in a second laboratory confirmed the ligand-dependent optical response and further defined its spectral properties. Upon UDP-Glc binding, absorption increased near 398 nm and decreased near 490 nm, consistent with a redistribution toward the neutral chromophore form (**Fig. 1E**). The corresponding extinction coefficients increased from 17.3 to 30.9 mM⁻¹ cm⁻¹ at 398 nm and decreased from 27.4 to 5.0 mM⁻¹ cm⁻¹ at 490 nm, whereas fluorescence quantum yields remained high, ranging from 0.59 to 0.74 across the two excitation wavelengths and ligand conditions (**Supplementary Table 1**). Together, these measurements support a substantial contribution of chromophore ionization changes to the ratiometric response.

Two-photon excitation brightness spectra likewise showed increased brightness near 780 nm and decreased brightness near 920 nm upon UDP-Glc binding, yielding an approximately 20-fold increase in the 780/920-nm excitation ratio (**Fig. 1F**). The wavelength-dependent brightness ratios highlight these opposing responses and demonstrate the potential for two-photon ratiometric imaging (**Fig. 1G**). At 940 nm, two-photon brightness decreased approximately fourfold, from 7.4 GM in the ligand-free state to 1.84 GM in the ligand-bound state (**Supplementary Table 1**). Together, these measurements establish that the pronounced ligand-dependent ratiometric response of UGluS extends to two-photon excitation.

We next evaluated metabolite responsiveness using a panel comprising UDP-Glc, UMP, UDP, UTP, glucose-1-phosphate, UTP plus glucose-1-phosphate, and related nucleotide sugars. UGluS responded preferentially to UDP-Glc, whereas the other tested analytes produced much smaller or negligible responses under the assay conditions (**Fig. 1H**). The control sensor dUGluS showed little response to the tested metabolites. Titration with UDP-Glc yielded an apparent dissociation constant of 32.5 ± 1.4 μM and a Hill coefficient of 1.4 ± 0.1 (**Fig. 1I**). Together, these results establish a large, preferential ratiometric response to UDP-Glc and support further evaluation of UGluS in living systems.

### UGluS reports nutrient- and enzyme-dependent changes in mammalian cells

UGluS expressed in HEK293T cells was diffusely distributed and exhibited robust fluorescence. Following 6 h of glucose deprivation, addition of 25 mM glucose induced a rapid increase in the fluorescence excitation ratio, reaching approximately 80% above baseline, with most of the initial response occurring within the first 10 min (**Fig. 2A and Supplementary Video 1**). dUGluS showed no comparable response. In glucose-containing medium (25 mM glucose), supplementation with 20 mM uridine produced a more gradual increase in the UGluS ratio, whereas dUGluS remained comparatively stable (**Fig. 2B and Supplementary Video 2**). These measurements distinguish rapid glucose-responsive changes from the slower response to uridine and demonstrate time-resolved monitoring of nutrient-dependent UDP-Glc availability.

**Figure 2.**
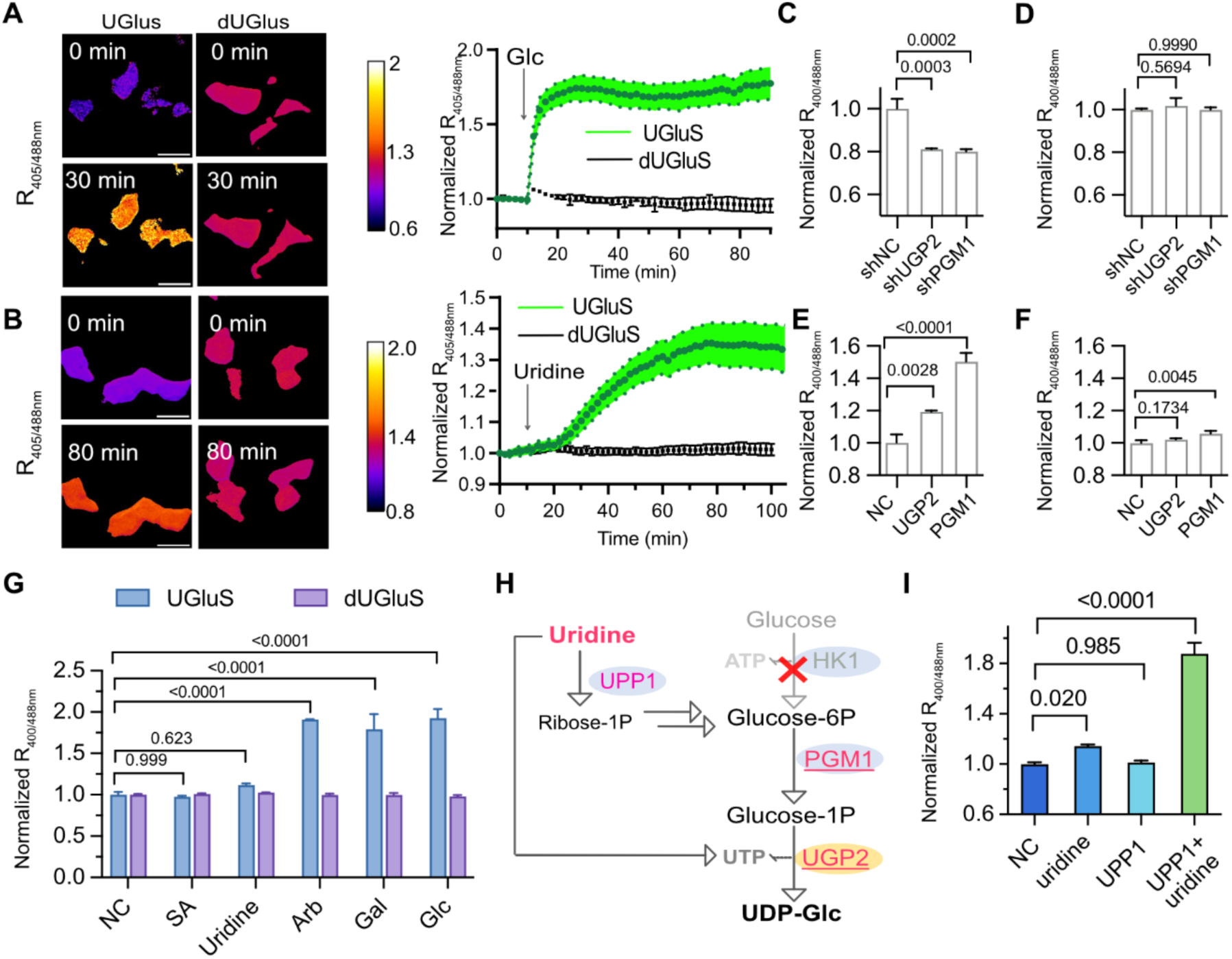
UGluS reports UDP-Glc responses to metabolic and genetic perturbations in HEK293T cells. (**A, B**) Representative ratiometric images and time-course traces of normalized fluorescence excitation ratios in HEK293T cells expressing UGluS or dUGluS following glucose (**A**; 25 mM) or uridine (**B**; 20 mM) supplementation. Scale bars, 20 μm. Panel **A** shows unnormalized fluorescence excitation-ratio images (*R* = *F*_405_/*F*_488_). In **B**, ratios were quantified over individual cells and normalized to the corresponding ratio at the start of imaging (*R*/*R*_0_, where *R*_0_ is the ratio at 0 h). (**C, D**) Responses of UGluS (**C**) and dUGluS (**D**) following shRNA-mediated knockdown of UGP2 or PGM1. shNC denotes a non-targeting control shRNA. (**E, F**) Responses of UGluS (**E**) and dUGluS (**F**) following overexpression of UGP2 or PGM1. NC, untreated negative control. **(G)** Responses of UGluS and dUGluS to metabolic precursors in glucose-starved cells. NC, negative control; SA, sialic acid; Ara, arabinose; Gal, galactose; Glc, glucose. **(H)** Schematic of the proposed contribution of uridine-derived carbon to UDP-Glc biosynthesis through UPP1 (uridine phosphorylase 1) under glucose-limited conditions. **(I)** UGluS responses to uridine supplementation, UPP1 overexpression, or their combination. Excitation ratios are 405/488 nm in **A** and **B** and 400/488 nm in **C**–**G** and **I**. Data in **A** and **B** are presented as mean ± SD (*n* = 12 cells pooled from 3 biological replicates per group). Data in **C**–**G** and **I** are presented as mean ± SD (*n* = 3 replicate wells per group). *P* values for the indicated comparisons in **C**–**G** and **I** were determined by one-way ANOVA with Dunnett’s multiple-comparisons test.

To test whether the sensor responds to genetic perturbations of UDP-Glc biosynthesis, we altered expression of UGP2 and phosphoglucomutase 1 (PGM1), two key enzymes in the biosynthetic pathway (**Fig. 1A and Supplementary Table 2**). shRNA-mediated knockdown of either enzyme reduced the UGluS fluorescence excitation ratio, with little change in dUGluS (**Fig. 2C,D**). Conversely, overexpression of UGP2 or PGM1 increased the UGluS ratio (**Fig. 2E**). The corresponding dUGluS responses were substantially smaller (**Fig. 2F**). Together, these bidirectional responses demonstrate that UGluS can report changes in intracellular UDP-Glc availability resulting from genetic manipulation of its biosynthetic pathway.

We next used UGluS to investigate the metabolic flexibility of the intracellular UDP-Glc pool. In glucose-deprived HEK293T cells, supplementation with arabinose or galactose markedly increased the UGluS fluorescence ratio to an extent comparable to glucose, whereas sialic acid had little effect (**Fig. 2G**). These findings suggest that alternative carbohydrate sources can support UDP-Glc replenishment under glucose-limited conditions. In contrast, uridine supplementation alone produced only a modest response. Because uridine phosphorylase 1 (UPP1) converts uridine into ribose-1-phosphate, which can subsequently enter central carbon metabolism,^30,31^ we investigated whether UPP1 expression influences the contribution of uridine to UDP-Glc biosynthesis (**Fig. 2H**). Neither uridine supplementation nor UPP1 overexpression alone substantially increased the UGluS ratio under these conditions. Strikingly, their combination produced a pronounced increase (**Fig. 2I**), consistent with enhanced channeling of uridine-derived carbon into UDP-Glc biosynthesis when UPP1 activity is elevated. These findings suggest that the contribution of uridine to UDP-Glc synthesis may depend on the cellular capacity for uridine catabolism and could become substantial in cell types, tissues, or physiological contexts with elevated UPP1 expression.^30,31^ Together, these results reveal the metabolic plasticity of the intracellular UDP-Glc pool and demonstrate how its availability can be dynamically regulated by distinct nutrient inputs and metabolic pathways.

### UGluS reports Yoda1-associated changes in cellular UDP-Glc

During live-cell assay optimization, we observed that routine cell handling could alter intracellular UGluS signals. To investigate this effect, HEK293T cells suspended in PBS were pipetted using either sharp or blunt tips to impose different degrees of mechanical perturbation. Cells handled with sharp tips exhibited lower UGluS fluorescence excitation ratios than those handled with blunt tips (**Supplementary Fig. 4A**). We next examined whether extracellular UDP-Glc could influence the sensor response following mechanical perturbation. Addition of extracellular UDP-Glc increased the UGluS ratio in sharp-tip-treated cells, and this response was attenuated by carbenoxolone (CBX), an inhibitor of connexin hemichannels and pannexin channels (**Supplementary Fig. 4B**).^32^ Given previous evidence that pannexin channels and connexin hemichannels mediate the passage of signaling molecules and metabolites,^16,33,34^ these findings suggest that CBX-sensitive, channel-associated membrane permeability may contribute to changes in intracellular UDP-Glc availability following mechanical perturbation.

Because pipetting could induce both mechanosensitive signaling and membrane damage, we next investigated the relationship between mechanosensitive signaling and intracellular UDP-Glc using pharmacological activation of Piezo1, a mechanosensitive Ca^2+^ channel activated by Yoda1.^35^ In HEK293T cells co-expressing UGluS and the red fluorescent Ca^2+^ indicator R-GECO, ^36^ addition of 30 μM Yoda1 elicited oscillatory Ca^2+^ signals accompanied by a gradual decrease in the UGluS fluorescence excitation ratio (**Fig. 3A,B and Supplementary Video 3**).

**Figure 3.**
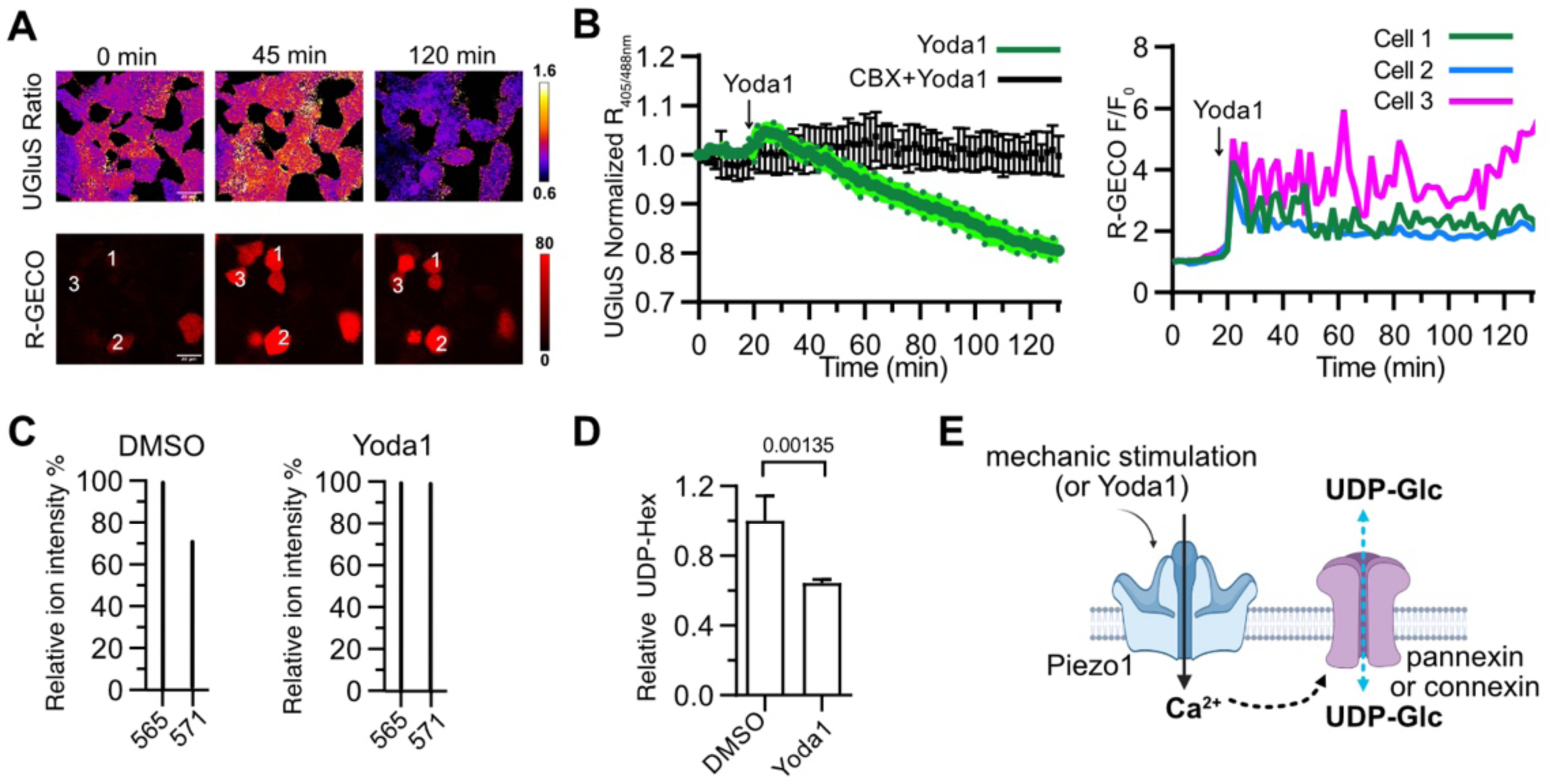
UGluS reveals Yoda1-induced changes in intracellular UDP-glucose and associated Ca^2+^ signaling. **(A)** Representative ratiometric images of UGluS (405/488 nm) and fluorescence images of R-GECO in HEK293T cells at 0, 45 and 120 min after the start of imaging, with 30 μM Yoda1 added at the time indicated in (B). Scale bars, 20 μm. **(B)** Time-course traces of normalized UGluS fluorescence excitation ratios (405/488 nm; left) in HEK293T cells following Yoda1 (30 μM) stimulation in the absence or presence of carbenoxolone (CBX, 50 μM). Ratios were normalized to their values at the start of imaging (0 min). Data are mean ± SD from 10 cells pooled from 3 biological replicates per group. Right, representative R-GECO fluorescence traces from three individual cells following Yoda1 stimulation, normalized to their initial fluorescence intensity (*F/F*_0_). Individual traces are shown because cell-to-cell variability and oscillatory Ca²⁺ responses complicate population averaging. **(C)** Representative selected-ion monitoring (SIM) signals at *m/z* 565 (endogenous UDP-hexose) and *m/z* 571 (added ^13^C_6_-UDP-glucose internal standard) from HILIC-MS analysis of extracts from DMSO-treated and Yoda1-treated HEK293T cells. **(D)** HILIC-MS quantification of relative intracellular UDP-hexose abundance in HEK293T cells treated with DMSO or 30 μM Yoda1. Data are mean ± SD (*n* = 3 biological replicates per group). The *P* value was determined using an unpaired, two-tailed Student’s *t*-test. **(E)** Proposed model of mechanosensitive regulation of intracellular UDP-Glc. Mechanical stimulation or pharmacological activation of Piezo1 by Yoda1 induces Ca^2^⁺ signaling, potentially promoting channel-mediated UDP-Glc release or redistribution.

Pretreatment with 50 μM CBX markedly attenuated the UGluS response (**Fig. 3B**). To obtain an orthogonal biochemical measurement, we analyzed extracts from DMSO- and Yoda1-treated HEK293T cells using hydrophilic interaction liquid chromatography–mass spectrometry (HILIC-MS). The endogenous UDP-hexose signal (*m/z* 565) coeluted with the ^13^C_6_-UDP-Glc internal standard (*m/z* 571) (**Supplementary Fig. 5**). Selected-ion monitoring (SIM) intensities measured within the coelution window were used to calculate endogenous UDP-hexose-to-internal-standard ratios. Consistent with the decrease reported by UGluS, Yoda1 treatment significantly reduced the relative intracellular UDP-hexose abundance (**Fig. 3C,D**).

Previous studies have demonstrated functional coupling between Piezo1 activation and connexin hemichannels or pannexin channels.^37,38^ Notably, PANX1-dependent release of UDP-Glc has been demonstrated in apoptotic cells,^16,33^ whereas connexin hemichannels have been shown to transport various metabolites, including ATP and glucose.^34^ Together, these findings and our fluorescence imaging and biochemical measurements support a model in which Piezo1 activation alters intracellular UDP-Glc availability through a CBX-sensitive process, potentially involving channel-mediated UDP-Glc release or redistribution (**Fig. 3E**).

### UGluS resolves UDP-glucose dynamics and regional differences in zebrafish

To evaluate UGluS performance in vivo, we generated zebrafish lines expressing UGluS or the unresponsive control dUGluS by Tol2-mediated transgenesis.^39^ Confocal time-lapse imaging of larvae at 5–7 days post-fertilization revealed that uridine supplementation robustly increased UGluS fluorescence excitation ratios, whereas dUGluS remained largely unchanged (**Fig. 4A,B and Supplementary Video 4**). These results demonstrate the ability of UGluS to monitor nutrient-induced UDP-Glc dynamics in real time in intact vertebrate organisms.

**Figure 4.**
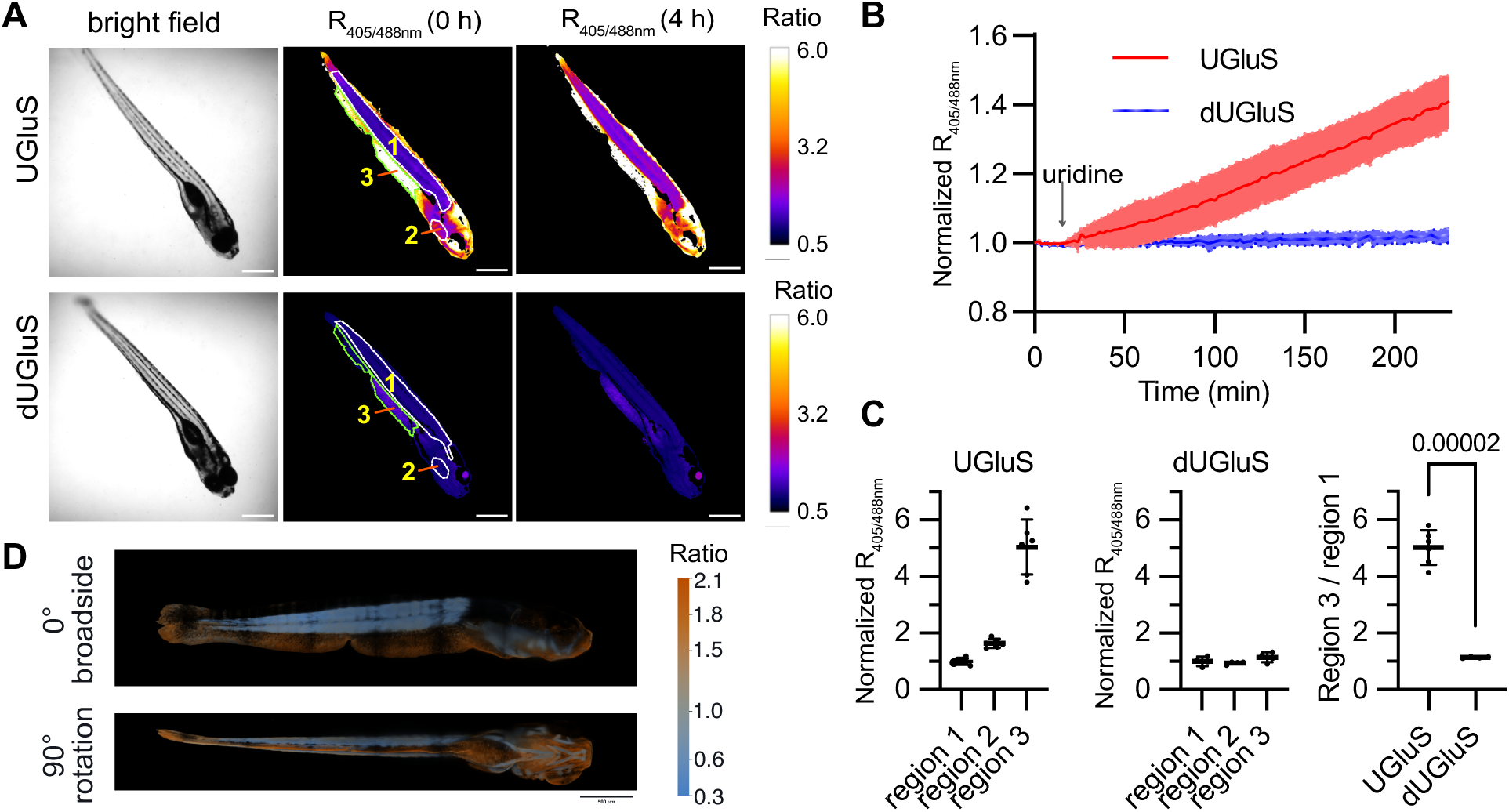
UGluS enables imaging of UDP-Glc dynamics and spatial heterogeneity in zebrafish larvae. (**A, B**) Representative images (**A**) and time courses of normalized fluorescence excitation ratios (405/488 nm; **B**) from zebrafish larvae (5–7 days post-fertilization) expressing UGluS or dUGluS following supplementation with 50 mM uridine. Panel **A** shows one bright-field image and unnormalized fluorescence excitation-ratio images (*R* = *F*_405_/*F*_488_) acquired at 0 and 4 h for each sensor. In **B**, ratios were quantified over the whole larva and normalized to the corresponding ratio at the start of imaging (*R*/*R*_0_, where *R*_0_ is the ratio at 0 h). Data in **B** are mean ± SD (*n* = 6 larvae for UGluS and *n* = 4 larvae for dUGluS). Scale bars, 100 μm. **(C)** Normalized fluorescence excitation ratios (405/488 nm) in regions 1–3 outlined in **A**, with region 3 corresponding to the ventral fin-fold region. Regional values were normalized to the mean region-1 ratio across larvae within each sensor group. The accompanying plot shows the region 3/region 1 quotient calculated separately for each larva. Points represent individual larvae, and summaries show mean ± SD (n = 6 larvae for UGluS and *n* = 4 larvae for dUGluS). Per-larva region 3/region 1 quotients were compared between sensor groups using a two-sided unpaired Welch’s *t*-test (*P* = 0.00002). (**D**) Representative light-sheet ratiometric images of UGluS-expressing zebrafish larvae shown in views separated by a 90° rotation. Scale bar, 500 μm. Similar results were obtained in two independent larvae. Color represents normalized excitation ratios, with brightness-modulated display as described in Methods.

Spatial analysis further revealed striking regional differences in UDP-Glc availability. UGluS fluorescence excitation ratios were markedly elevated in the ventral fin fold (region 3; **Fig. 4A**) compared with other regions of the larval body, whereas dUGluS exhibited substantially smaller regional differences (**Fig. 4C**). Importantly, the relative enrichment in region 3 compared with region 1 was significantly greater for UGluS than for dUGluS (**Fig. 4C**), supporting a sensor-specific component of the observed spatial heterogeneity. Light-sheet ratiometric imaging further visualized the heterogeneous distribution of UGluS signals in intact larvae (**Fig. 4D and Supplementary Video 5**), providing complementary evidence for spatially distinct UDP-Glc availability. The fin fold undergoes active morphogenesis and extracellular matrix remodeling,^40–42^ processes that require substantial biosynthetic activity and may contribute to the elevated UDP-Glc availability observed in this region. Together, these findings demonstrate that UGluS enables real-time imaging of metabolic responses and reveals previously inaccessible spatial heterogeneity in UDP-Glc availability during vertebrate development.

### UGluS reveals dynamic regulation of UDP-Glc in mouse tissues ex vivo and in vivo

To extend UGluS imaging to mammalian tissues, we expressed UGluS or dUGluS in mouse liver using adeno-associated virus (AAV) vectors driven by the liver-specific thyroxine-binding globulin promoter.^43^ Fresh liver slices were subsequently prepared for confocal imaging (**Fig. 5A**). Insulin stimulation induced a pronounced, time-dependent decrease in the UGluS fluorescence excitation ratio, whereas dUGluS remained largely unchanged (**Fig. 5B–D**). Notably, insulin treatment also markedly increased hepatic glycogen content (**Fig. 5E**). Because UDP-Glc serves as the immediate glucose donor for glycogen synthesis,^44,45^ these findings reveal that insulin-stimulated glycogen accumulation is accompanied by a substantial reduction in intracellular UDP-Glc availability, consistent with enhanced utilization of the metabolite during glycogenesis.

**Figure 5.**
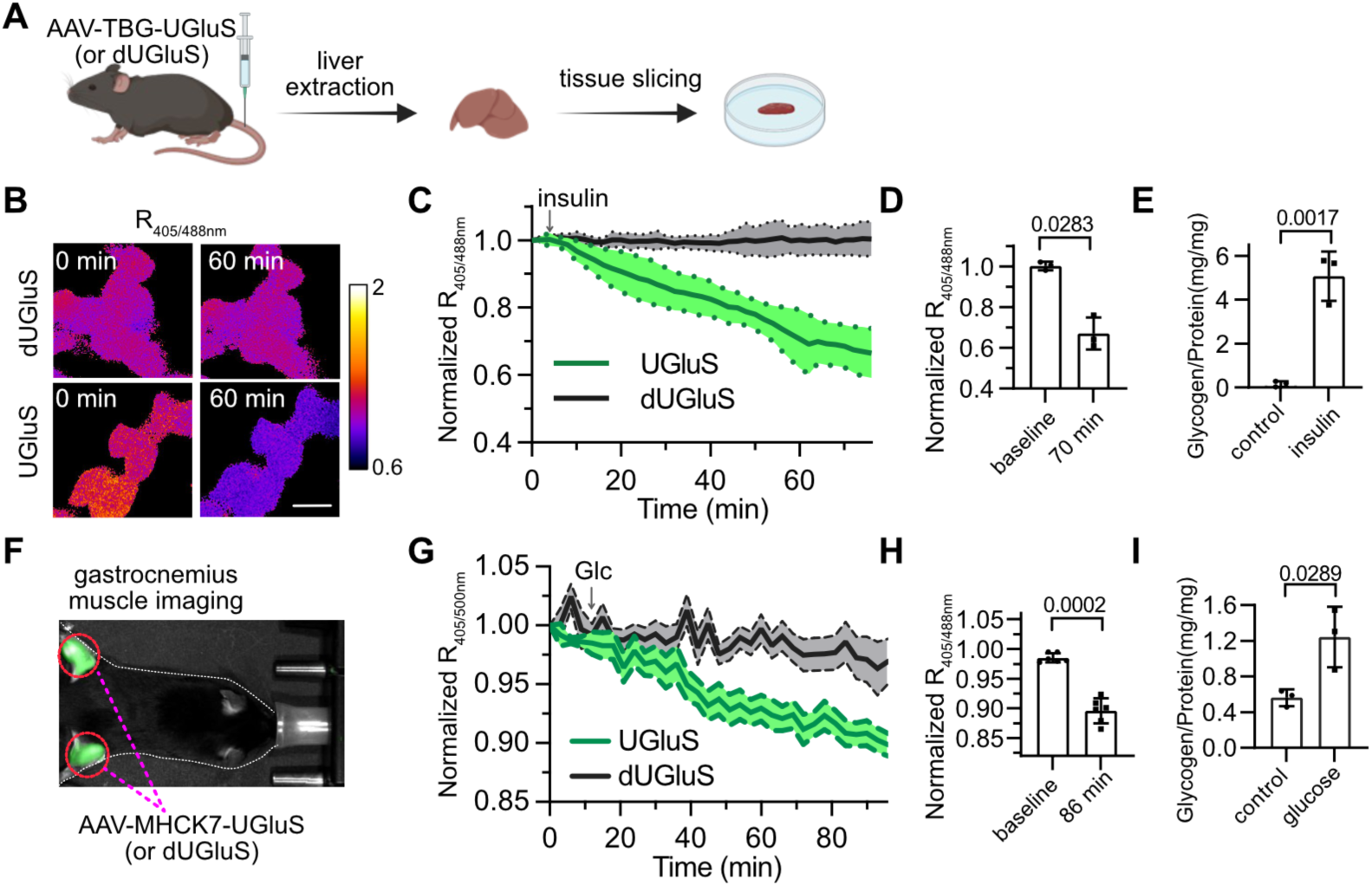
UGluS enables imaging of UDP-Glc dynamics in mouse tissues ex vivo and in vivo. (**A**) Schematic of ex vivo liver imaging. Mice received intravenous injections of adeno-associated virus (AAV) vectors expressing UGluS or dUGluS under the control of the TBG promoter. Fresh liver slices were subsequently prepared for imaging. (**B, C**) Representative ratiometric images acquired at 0 and 60 min (**B**) and time courses of normalized fluorescence excitation ratios (405/488 nm; **C**) in liver slices expressing UGluS or dUGluS following treatment with 500 nM insulin. Fluorescence excitation ratios were normalized to their initial values at the start of imaging (*R/R*_0_). Data in **C** are mean ± SD (*n* = 3 liver slices from 3 independent mice per group). Scale bar, 20 μm. **(D)** Quantification of normalized UGluS fluorescence excitation ratios immediately before insulin treatment (baseline) and at 70 min after treatment. Each point represents a mouse-level measurement, with the same mice analyzed at both time points. Data are mean ± SD (*n* = 3 mice). **(E)** Glycogen content normalized to total protein content in liver slices treated with Krebs– Henseleit buffer (control) or insulin. Data are mean ± SD (*n* = 3 liver slices from 3 mice per group). **(F)** Representative setup for in vivo ratiometric imaging of gastrocnemius muscle expressing UGluS or dUGluS delivered by AAV vectors under the control of the MHCK7 promoter. **(G)** Time courses of normalized fluorescence excitation ratios (405/500 nm; emission collected at 550 nm) in the gastrocnemius muscles of live mice expressing UGluS or dUGluS following intraperitoneal glucose administration (1.5 g/kg). Fluorescence excitation ratios were normalized to their initial values at the start of imaging (*R/R*₀). Measurements from the left and right gastrocnemius muscles were averaged for each mouse. Data are mean ± SEM (*n* = 6 mice per group). **(H)** Quantification of normalized UGluS fluorescence excitation ratios immediately before glucose treatment (baseline) and at 86 min after treatment. Each point represents the average of measurements from the left and right gastrocnemius muscles of one mouse (*n* = 6 mice). The same mice were analyzed at both time points. Data are mean ± SD. **(I)** Glycogen content normalized to total protein content in gastrocnemius muscle samples collected from mice following intraperitoneal administration of saline (control) or glucose. Data are mean ± SD (*n* = 3 mice per group). *P* values in **D** and **H** were determined using paired, two-tailed *t*-tests, while *P* values in **E and I** were determined using unpaired, two-tailed *t*-tests.

We next investigated whether UGluS could resolve UDP-Glc dynamics in living mice. UGluS or dUGluS was expressed in gastrocnemius muscle using AAV vectors driven by the muscle-specific MHCK7 promoter (**Fig. 5F**).^46^ Following intraperitoneal glucose administration in fasted mice, UGluS exhibited a progressive decrease in its fluorescence excitation ratio, whereas dUGluS showed a substantially smaller response (**Fig. 5G,H**). Consistent with the role of skeletal muscle in glucose disposal and glycogen storage,^47–49^ glucose administration also increased muscle glycogen content (**Fig. 5I**). These findings demonstrate that UGluS can monitor dynamic changes in UDP-Glc availability in vivo and reveal a coordinated increase in glycogen accumulation and decrease in the intracellular UDP-Glc pool following glucose administration.

Together, the liver and skeletal muscle experiments establish the utility of UGluS for imaging UDP-Glc dynamics in mammalian tissues ex vivo and in living animals. Importantly, both models reveal that stimulation of glycogen accumulation is accompanied by a decrease, rather than an increase, in UDP-Glc availability. These observations highlight the dynamic balance between UDP-Glc production and consumption and demonstrate the value of real-time imaging for uncovering metabolic responses that cannot be inferred from nutrient availability or glycogen accumulation alone.

## Discussion

Here, we developed UGluS, a genetically encoded fluorescent sensor that enables real-time visualization of UDP-Glc dynamics across cellular, tissue, and whole-animal systems. By repurposing UGP, an enzyme responsible for UDP-Glc biosynthesis, we created a sensor that combines an exceptionally large ratiometric response, preferential selectivity for UDP-Glc, and pronounced ligand-dependent optical responses under both one- and two-photon excitation.

Applications in cultured mammalian cells, developing zebrafish, mouse liver slices, and living mice reveal dynamic and spatially heterogeneous regulation of UDP-Glc under diverse metabolic and physiological conditions. Together, these advances establish UGluS as a powerful tool for investigating nucleotide-sugar metabolism and expand the range of enzyme-derived scaffolds available for genetically encoded metabolite sensing.

Enzymes represent a rich source of molecular-recognition scaffolds, yet exploiting the recognition of a biosynthetic product by the enzyme responsible for its synthesis remains a relatively underexplored sensor-engineering strategy. Building on previous enzyme-derived sensors, UGluS demonstrates the potential of product-directed sensing for nucleotide sugars. Through catalytic inactivation, structure-guided cpGFP insertion, and iterative directed evolution, we converted UGP into a highly responsive fluorescent sensor while preserving preferential recognition of UDP-Glc over related nucleotide sugars and reaction components. The approximately 18-fold ratiometric response, together with substantial ligand-dependent changes in two-photon excitation properties, highlights the optical performance achievable through this approach. Because biosynthetic enzymes collectively recognize a vast diversity of cellular metabolites, expanding the use of their product-binding properties could provide new opportunities for engineering fluorescent sensors targeting metabolites that remain difficult to monitor in living systems.

Beyond sensor development, UGluS reveals substantial metabolic flexibility in the regulation of intracellular UDP-Glc. In mammalian cells, glucose replenishment rapidly increased UDP-Glc availability, whereas uridine elicited a more gradual response. Genetic manipulation of UGP2 and PGM1 further demonstrated the sensitivity of the UDP-Glc pool to its biosynthetic machinery. Particularly intriguing was the pronounced increase in UGluS signal following uridine supplementation in cells overexpressing UPP1 under glucose-limited conditions. Recent studies have established that UPP1-mediated uridine catabolism provides an alternative carbon source under glucose restriction, supporting central carbon metabolism, cancer cell survival, and the biosynthesis of nucleotide sugars such as UDP-GlcNAc.^30,31^ Our findings extend this metabolic connection to UDP-Glc, suggesting that uridine-derived ribose may contribute to its replenishment when glucose is scarce. The dependence on UPP1 expression further suggests that this alternative metabolic route may become particularly important in cells with elevated UPP1 activity. Together, these findings highlight the metabolic plasticity of nucleotide-sugar homeostasis and demonstrate the utility of UGluS for uncovering nutrient-dependent metabolic adaptations.

The ability to image UDP-Glc in intact animals further extends these findings to spatially organized metabolic regulation. In zebrafish larvae, UGluS detected robust responses to uridine supplementation and revealed striking regional differences in UDP-Glc availability. In particular, the ventral fin fold exhibited substantially elevated UGluS fluorescence ratios, with significantly greater regional enrichment than observed with the unresponsive control dUGluS. Complementary light-sheet imaging further visualized the heterogeneous distribution of sensor signals. The fin fold undergoes active morphogenesis and extracellular matrix remodeling, processes requiring coordinated biosynthetic activity.^40–42^ Elevated UDP-Glc availability in this region may therefore reflect distinct metabolic demands associated with tissue development and remodeling. These findings highlight the potential of UGluS to investigate the spatial organization of nucleotide-sugar metabolism during development and other physiological processes, when metabolite distributions cannot be adequately captured by conventional bulk measurements.

The mouse experiments reveal a particularly interesting relationship between UDP-Glc availability and glycogen metabolism. In liver slices, insulin stimulation induced a pronounced decrease in the UGluS ratio, accompanied by increased glycogen accumulation. Similarly, glucose administration to living mice produced a measurable decrease in the UGluS ratio in skeletal muscle while increasing muscle glycogen content. These observations are notable because enhanced nutrient availability or anabolic stimulation might be expected to increase intracellular UDP-Glc production. Instead, the observed decreases suggest that UDP-Glc utilization during glycogen synthesis can exceed its replenishment under these conditions. Thus, intracellular UDP-Glc availability reflects the dynamic balance between metabolite production and consumption, rather than nutrient supply alone. Importantly, the ability of UGluS to resolve these responses in living tissues and animals provides an opportunity to investigate the regulation of nucleotide-sugar metabolism during physiological transitions, including nutrient uptake, glycogen synthesis, and metabolic adaptation.

Our findings also reveal a potentially important connection between mechanosensitive signaling and intracellular nucleotide-sugar homeostasis. Both mechanical perturbation and pharmacological activation of Piezo1 altered UGluS responses, with the Yoda1-induced decrease substantially attenuated by CBX. Independent HILIC-MS measurements further demonstrated reduced intracellular UDP-hexose abundance following Yoda1 treatment, providing biochemical support for the fluorescence observations. Previous studies have established PANX1-dependent UDP-Glc release from apoptotic cells and functional coupling between Piezo1 activation and connexin hemichannels or pannexin channels.^16,33,37,38^ Together, these findings support a model in which mechanosensitive signaling alters intracellular UDP-Glc availability through a CBX-sensitive process, potentially involving channel-mediated metabolite release or redistribution. Given the established roles of extracellular UDP-sugars as signaling molecules,^15–18^ this UGluS-informed model suggests a potential connection between cellular mechanosensing, intracellular nucleotide-sugar metabolism, and extracellular signaling.

Several considerations highlight opportunities for further development and application of UGluS. As with other genetically encoded metabolite sensors, UGluS reports dynamic changes in the sensor-accessible UDP-Glc pool rather than directly measuring metabolic flux. It therefore complements, rather than replaces, approaches such as isotope tracing and quantitative mass spectrometry, which provide information on metabolic pathways, carbon utilization, and metabolite abundance. Although our mechanosensitive experiments implicate a CBX-sensitive process, identifying the responsible channels and determining whether UDP-Glc release contributes to extracellular signaling will require targeted genetic perturbations and functional characterization of downstream signaling responses. Extending UGluS to monitor extracellular UDP-Glc dynamics may also require higher-affinity variants capable of detecting the lower metabolite concentrations expected in the extracellular environment. Further optimization of sensor affinity and subcellular targeting could broaden its applications to different cellular compartments and physiological contexts.

In summary, UGluS provides a versatile tool for visualizing UDP-Glc dynamics in living systems and reveals previously difficult-to-access aspects of nucleotide-sugar regulation, including nutrient-dependent metabolic flexibility, spatial heterogeneity during development, dynamic responses associated with glycogen synthesis, and potential coupling to mechanosensitive signaling. By combining a biosynthetic enzyme-derived recognition scaffold with high-performance ratiometric fluorescence and applications across diverse biological systems, this work advances genetically encoded nucleotide-sugar sensing and opens new opportunities for investigating the roles of UDP-Glc in metabolism, physiology, and disease.

## Methods

### Bacterial strains and mammalian cell culture

DH10B *Escherichia coli* (Thermo Fisher Scientific, 18297010) was used for cloning, protein engineering, and protein expression. NEB Stable *E. coli* (New England Biolabs, C3040) was used to propagate lentiviral and AAV transfer and packaging plasmids. Bacteria were cultured under standard conditions unless otherwise noted.

HEK293T cells were obtained from ATCC (CRL-3216) and used at passages below 25. Cells were maintained in high-glucose DMEM (Corning) supplemented with 10% fetal bovine serum, 100 U/mL penicillin, and 100 μg/mL streptomycin at 37 °C with 5% CO_2_. Sex was not considered as a biological variable in the immortalized cell-line experiments.

### Zebrafish and mouse husbandry and ethics

The xanthophore-deficient zebrafish *fms1*^j4e^^1^ line from the Parichy laboratory was used to reduce background tissue autofluorescence.^50^ Zebrafish were maintained at approximately 28 °C on a 14 h light/10 h dark cycle. Larvae were initially fed marine rotifers enriched with Algamac (Reed Mariculture), and older larvae and adults were fed live brine shrimp and GEMMA Micro 500 (Skretting). Procedures followed the NIH Guide for the Care and Use of Laboratory Animals and were approved by the University of Virginia Animal Care and Use Committee (protocol 4170). Fish were euthanized by MS-222 overdose followed by physical maceration. Sex was not determined for larval zebrafish, and sex-specific effects were not assessed.

Mouse procedures were approved by the University of Virginia Animal Care and Use Committee (protocol 4196). Female and male C57BL/6J mice (The Jackson Laboratory, strain 000664), 6–8 weeks old, were housed in a temperature- and humidity-controlled facility on a 12 h light/dark cycle, with chow and water available ad libitum unless otherwise indicated. Both sexes were included, but the study was not powered to detect sex-specific differences.

### Sources and construction of plasmids

pMD2.G (Addgene 12259) and psPAX2 (Addgene 12260) were gifts from Didier Trono; pLKO.1-TRC (Addgene 10878)^51^ was a gift from David Root; pAdDeltaF6 (Addgene 112867) and pAAV2/9n (Addgene 112865) were gifts from James M. Wilson; pWPI-UPP1-3xFLAG (Addgene 201644)^30^ was a gift from Alexis Jourdain and Vamsi Mootha; and CMV-R-GECO1 (Addgene 32444)^36^ was a gift from Robert Campbell. Restriction endonucleases and Phusion High-Fidelity DNA Polymerase were purchased from Thermo Fisher Scientific. Vector backbones were linearized by restriction digestion or PCR, and inserts were amplified with overlapping primers. DNA fragments were purified by agarose gel electrophoresis and gel extraction, assembled using an in-house Gibson assembly mixture, and used to transform electrocompetent DH10B cells. Clones were screened by colony PCR or restriction digestion and confirmed by Sanger or whole-plasmid sequencing through Eurofins Genomics.

Oligonucleotides are listed in Supplementary Table 3.

### Sensor engineering

The *H. pylori* UGP gene was synthesized by IDT and cloned into pBAD by Gibson assembly. The R15A substitution was introduced as described in the Results. The scaffold was amplified and linearized at the selected insertion sites, and a cpGFP fragment was inserted by Gibson assembly. NNK codons (N = A, T, G, or C; K = G or T) at both ends of the cpGFP fragment diversified the two fusion linkers in the insertion-site libraries. The subsequent linker-optimization libraries varied linker length and sequence. Library architectures are shown in Supplementary Fig. 1C, and the primer sequences and library-specific degenerate bases are listed in Supplementary Table 3.

Fluorescent colonies were inoculated into 96-deep-well plates containing 1 mL 2×YT medium with 0.01% L-arabinose and 100 μg/mL ampicillin per well and grown overnight at 37 °C with shaking at 250 rpm. Cells were lysed in 200 μL buffer containing 5 mg/mL octyl glucoside, 0.1 mg/mL chicken egg lysozyme, and 0.2 U/mL Benzonase in 20 mM Tris-HCl, pH 8.0, for 1 h at 16 °C with shaking at 250 rpm. After addition of 600 μL 50 mM Tris-HCl, pH 7.4, plates were centrifuged at 3,000 × *g* for 10 min. Aliquots of clarified lysate (50 μL) were transferred to paired 96-well assay plates. One plate received 50 μL 100 μM UDP-Glc in 50 mM Tris-HCl, pH 7.4, and the control plate received 50 μL buffer. Endpoint fluorescence was measured using a BioTek Synergy Mx microplate reader. Each screened colony contributed one measurement per condition, without technical replication.

Variants with the largest changes in the 400/488-nm fluorescence excitation ratio were advanced. Following insertion-site and linker optimization, random mutagenesis was performed by error-prone PCR using a previously described procedure.^7^ Brightening mutations were introduced into UGluS0.2 by multisite mutagenesis and Gibson assembly with mutagenic oligonucleotides. The evolution sequence and final mutations are shown in Supplementary Figs. 1–3.

### Protein purification and fluorescence assays

A single colony of DH10B cells carrying pBAD-UGluS or pBAD-dUGluS was grown overnight in 2 mL 2×YT with 100 μg/mL ampicillin at 37 °C and 250 rpm. The starter culture was used to inoculate 500 mL 2×YT containing 0.02% L-arabinose and 100 μg/mL ampicillin. Cultures were grown for 2–5 h at 37 °C and then for 48 h at 16 °C. Cells were collected at 5,000 × *g* for 20 min and resuspended in 30 mL lysis buffer containing 50 mM sodium phosphate, 300 mM NaCl, and 10 mM imidazole, pH 7.4, with freshly added 1 mM PMSF. Following sonication, lysates were clarified at 10,000 × *g* for 30 min at 4 °C using a JA-20 rotor (Beckman Coulter). His_6_-tagged proteins were purified by Ni-NTA chromatography and size-exclusion chromatography on a HiLoad 16/600 Superdex 200 pg column (Cytiva) in 50 mM Tris-HCl, pH 7.4.

Purified protein was diluted to approximately 100 nM in 50 mM Tris-HCl, pH 7.4. Spectral measurements used protein with 100 μM UDP-Glc or buffer alone. Excitation spectra were recorded from 350 to 500 nm with emission at 520 nm, and emission spectra from 500 to 650 nm with excitation at 480 nm. Endpoint assays used excitation at 400 and 488 nm and emission at 520 nm. Specificity assays tested 100 μM UMP, UDP, UTP, glucose-1-phosphate, UTP plus glucose-1-phosphate, UDP-GlcNAc, UDP-Gal, UDP-GalNAc, GDP-Fuc, or GDP-Man. UDP-Glc titrations covered 5–200 μM, and responses were fitted by nonlinear regression to the Hill equation to estimate the apparent dissociation constant and Hill coefficient.

### Photophysical characterization

One-photon absorption spectra were measured using a Lambda 950 spectrophotometer (PerkinElmer). The extinction coefficient of the anionic chromophore in the UDP-Glc-saturated state was determined using the Strickler–Berg method.^52^ Extinction coefficients of the neutral chromophore in free and saturated states were calculated using the effective extinctions of wild-type GFP neutral and anionic forms (ρ_N_ε_N_ = 30,000 M^−1^ cm^−1^ and ρ_A_ε_A_ = 7,000 M^−1^ cm^−1^, respectively; ρ_N_ + ρ_A_ = 1) and an EGFP anionic extinction coefficient of 55,900 M^−1^ cm^−1^.

Fractional populations of the sensor chromophore forms were obtained from the optical densities of their absorption peaks and the extinction coefficients.^53^ The fluorescence quantum yield of the anionic form was measured with a Quantaurus-QY integrating-sphere spectrofluorimeter (Hamamatsu). Fluorescence lifetimes were measured with a ChronosDFD digital frequency-domain system coupled to a PC1 spectrofluorometer (ISS). Two-photon excitation spectra and cross sections of the free and saturated states were measured as previously described.^54^

### Cellular metabolic and genetic perturbations

For glucose re-addition experiments, HEK293T cells expressing UGluS or dUGluS were deprived of glucose for 6 h before stimulation with 25 mM glucose. For the uridine time-course experiments, 20 mM uridine was added under glucose-containing conditions (25 mM glucose). For supplementation experiments, glucose-deprived cells were treated with 20 mM uridine, arabinose, galactose, sialic acid, or glucose for 2 h. UGP2 and PGM1 were perturbed by shRNA-mediated knockdown or overexpression, and UPP1 was tested by overexpression with or without uridine. For shRNA stably expression cell populations, HEK293T cells in 6-well plates were transduced at approximately 50% confluence with 300 μL filtered lentiviral supernatant per well. After 48 h, cells were passaged and selected for 1 week with 1 μg/mL puromycin. shRNA target sequences are listed in Supplementary Table 2, and cloning oligonucleotides are listed in Supplementary Table 3.

For the mechanical-handling assay, cells suspended in PBS were pipetted using sharp or blunt 200-μL tips. Sharp-tip-treated cells were subsequently exposed to extracellular UDP-Glc (500 μM) with or without CBX. For Yoda1 experiments, cells co-expressed UGluS and R-GECO and were stimulated with 30 μM Yoda1. CBX pretreatment was performed at 50 μM for 20 min at 37 °C before imaging and Yoda1 addition.

### Fluorescence microscopy of mammalian cells

Cells were seeded onto 7 × 7 mm glass coverslips in 24-well plates. Imaging used an inverted Leica DMi8 microscope with an SPE-II spectral confocal module. Before imaging, coverslips were transferred to a 35-mm glass-bottom dish containing 2 mL HBSS supplemented with 20 mM HEPES, pH 7.4, and equilibrated at room temperature for approximately 1 h. Images were collected with a 40× oil-immersion objective (NA 1.3), sequential 405- and 488-nm excitation at approximately 5% laser power, and emission at 500–550 nm. Time-lapse image sets were acquired every 2 min.

Yoda1 experiments were performed at 37 °C in phenol red-free DMEM (Thermo Fisher Scientific, A1443001) supplemented with 4.5 mg/mL glucose, 20 mM HEPES, pH 7.4, 1 mM sodium pyruvate, and 1× GlutaMax (Thermo Fisher Scientific, 35050061). For inhibition experiments, CBX was applied as described above. R-GECO fluorescence was imaged sequentially after the two imaging channels of UGluS, using 532-nm laser excitation at approximately 8% laser power, with emission collected from 570 to 650 nm.

### Plate-reader measurements of cellular fluorescence

Cells in 24-well plates were subjected to the indicated treatments and, near confluence, washed three times with 200 μL DPBS containing Ca^2+^ and Mg^2+^. Cells were resuspended in 200 μL DPBS and transferred to 96-well plates. Fluorescence was measured using a BioTek Synergy Mx with excitation at 400 and 488 nm, emission at 520 nm, a 9-nm bandwidth, and a detector gain of approximately 100. Excitation ratios were calculated from the corresponding fluorescence signals.

### Zebrafish transgenesis and imaging

pDestTol2-ubic-UGluS and pDestTol2-ubic-dUGluS were assembled using the ubi promoter, Tol2Kit destination vectors, and the corresponding sensor coding sequences.^39^ Plasmids were injected into one-cell embryos together with Tol2 transposase mRNA (25 pg per embryo).^55^ Embryos were screened for fluorescence at 3 days post-fertilization. Stable founders were identified by outcrossing, and F3 larvae at 5–7 days post-fertilization were used for imaging.

For confocal imaging, larvae were embedded in 2% low-melting-point agarose prepared in 1× Hank’s solution containing 0.008% tricaine at approximately 37–40 °C in 3.5-cm dishes. Following solidification, larvae were imaged on a Leica DMi8 with a 5× objective. Sequential 405- and 488-nm excitation was performed at approximately 5% laser power, with emission at 500–550 nm. Images were acquired at 2-min intervals. Uridine was added at 50 mM for the time-course experiments. Whole-larva excitation ratios were normalized to the corresponding ratio at the start of imaging (R/R_0_); six UGluS-expressing and four dUGluS-expressing larvae were analyzed.

For regional analysis, fluorescence excitation ratios were quantified in regions 1–3 outlined in Fig. 4A, with region 3 corresponding to the ventral fin-fold region. For display, values for all three regions were divided by the mean region-1 ratio across larvae within the same sensor group. Separately, a region 3/region 1 quotient was calculated from the two matched regional measurements for each individual larva and used for the between-sensor comparison.

For light-sheet imaging, larvae were embedded as above and drawn into a capillary before the agarose solidified. The capillary was mounted, and a small agarose segment was extruded to position the larva in the imaging chamber. The chamber contained 1× Hank’s solution with 0.008% tricaine. Imaging used a Zeiss Lightsheet 7 with a W Plan-Apochromat 20×/1.0 Corr = 1.33 objective at 0.7× zoom. Voxel size was 0.33 × 0.33 × 1 μm. Sequential excitation used 405 and 488 nm at 5% laser power; emission was collected at 505–545 nm. Exposure was 30 ms without binning.

### Lentiviral and AAV production

For lentivirus production, HEK293T cells at approximately 70% confluence in 6-well plates were transfected with 1,000 ng transfer plasmid, 1,000 ng psPAX2, and 500 ng pMD2.G in 200 μL Opti-MEM using 7.5 μL of 1 μg/μL linear PEI (MilliporeSigma, 764965). Supernatants were collected after 48 h, filtered through 0.45-μm filters, and stored at −80 °C.

AAV production used pAdDeltaF6, pAAV2/9n, and one of pAAV-MHCK7-UGluS, pAAV-MHCK7-dUGluS, pAAV-TBG-UGluS, or pAAV-TBG-dUGluS. HEK293T cells were co-transfected and maintained for 96 h in high-glucose DMEM containing 4% fetal bovine serum, 0.1 M sucrose, and 10 mM HEPES. Cells and conditioned medium were collected for purification by gradient ultracentrifugation. Virus was buffer-exchanged into DPBS containing 0.001% Pluronic and 200 mM NaCl, aliquoted, flash-frozen, and stored at −80 °C. Titers were measured by quantitative PCR according to the Addgene protocol and were typically approximately 1 × 10^13^ genome copies/mL.

### Mouse liver slice preparation and imaging

C57BL/6J mice received retro-orbital injections of AAV encoding TBG-driven UGluS or dUGluS (150 μL total, 75 μL per eye). Imaging was performed 3–5 weeks later. Following a 4 h fast, mice were anesthetized with ketamine (100 mg/kg) and xylazine (10 mg/kg), administered intraperitoneally. Livers were excised and placed in ice-cold, carbogenated (95% O_2_/5% CO_2_) modified Krebs–Henseleit buffer (KHB) containing 118.0 mM NaCl, 4.7 mM KCl, 2.52 mM CaCl_2_, 1.64 mM MgSO_4_, 24.88 mM NaHCO_3_, 1.18 mM KH_2_PO_4_, and 5.55 mM glucose, pH 7.4, 298 mOsm/kg. Liver lobes were sectioned by vibratome in ice-cold KHB. The superficial 250 μm containing Glisson’s capsule was discarded, and experimental slices were cut at 150 μm.

Slices were transferred to recovery KHB at 37 °C containing 1 mM sodium pyruvate, 2 mM L-glutamine, 0.3 mM glycine, 0.4 mM cysteine, and 10 mM HEPES, pH 7.4, and recovered for 1 h with continuous carbogen bubbling. Slices were then imaged in recovery buffer in 35-mm dishes on a Leica DMi8 confocal microscope at 37 °C with continuous carbogen perfusion. Images were acquired every 2 min using a 40× oil-immersion objective (NA 1.3), sequential 405- and 488-nm excitation at approximately 5% laser power, and emission at 500–550 nm. Following a short baseline, insulin was added to 500 nM, and imaging continued for about 90 min.

### In vivo imaging of mouse gastrocnemius muscle

AAV vectors driven by MHCK7 were injected bilaterally into gastrocnemius muscles (50 μL total), and expression proceeded for 3 weeks. Mice were fasted for 4 h and anesthetized with isoflurane (5% induction, 1.5% maintenance). Hair overlying the hindlimbs was shaved and depilated. Imaging used a Lago X system with a thermoelectrically cooled (−90 °C) Model 1900 CCD camera (Spectral Instruments Imaging; Aura 4.5.1). Sequential LED excitation used 405 and 500 nm at 30% power, and emission was collected at 550 nm. Fluorescence acquisition used a 10-s exposure, f/2.0 aperture, detector gain 0.51, 2×2 binning, and a 25-cm field of view.

Brightfield images used the same field of view and binning, a 0.5-s exposure, and f/16.0 aperture. Hardware flat-field and dark-noise corrections were applied, and acquisition parameters were held constant across replicates. Imaging comprised 40 sequential iterations. After seven baseline iterations, glucose (1.5 g/kg) was administered intraperitoneally, and imaging continued to follow the sensor response. Normalized fluorescence excitation ratios from the left and right gastrocnemius muscles were averaged within each mouse at each time point; the mouse was the independent biological replicate (n = 6 per group).

### Glycogen content measurement

Glycogen levels in liver and skeletal muscle tissues (∼20 mg) were quantified using a commercial Colorimetric Glycogen Assay Kit (ab65620, Abcam). Tissues were homogenized in ice-cold ddH_2_O (200 μL), boiled for 10 min to inactivate endogenous enzymes, and centrifuged at 18,000 × g for 10 min (4 °C) to harvest the supernatant. Aliquots of the supernatant were incubated with Hydrolysis Enzyme Mix I for 30 min at room temperature to hydrolyze glycogen into glucose. Samples were subsequently reacted with the colorimetric reaction mix containing OxiRed probe for 30 min in the dark, and absorbance was measured at 570 nm using a microplate reader. Sample background controls without hydrolysis enzyme were included to subtract baseline glucose interference. Concentrations were determined using a glycogen standard curve and normalized to total protein content.

### Image processing

Ratiometric images were processed using Fiji^56^ following a reported procedure.^57^ Images from the two excitation channels underwent rolling-ball background subtraction with a 50-pixel radius. For the 405/488-nm microscopy datasets, the 405-nm excitation stack was aligned to the 488-nm excitation stack using a rigid-body transformation in MultiStackReg. Images were converted to 32-bit format and smoothed to reduce high-frequency noise. An intensity threshold masked background pixels outside the region of interest as Not a Number (NaN). Ratio Plus was used to divide the 405-nm excitation image by the corresponding 488-nm excitation image. Ratio images were exported as TIFF files for pseudocolor display and analysis.

To process the light-sheet microscopy data, images were exported from Imaris and imported into Fiji plane by plane. A background offset of 88 intensity counts, estimated from dim regions in both channels, was subtracted from each channel. For each pixel, R_405/488_ was calculated as (I_405_-88)/(I_488_-88). Pixels with corrected 488-nm intensity below 25 counts were masked in both the ratio and brightness images. Corrected brightness was calculated as [(I_405_-88)+(I_488_-88)]/2. The ratios were divided by a single reference value, 0.4715962287, calculated as the pooled arithmetic mean of all valid ratio pixels from five randomly selected middle slices. The same normalization factor was applied to all 689 sections. Next, valid normalized ratios and corrected brightness values were averaged separately within 3×3 XY bins, retaining all 689 Z planes. The resulting voxel spacing was approximately 0.99 × 0.99 × 1.00 µm. The mean ratio determined the color using a blue-to-neutral-to-orange lookup table spanning 0.3–2.1. Ratios outside that interval were assigned the nearest endpoint color. Corrected brightness was mapped linearly over 0–4,500 counts and used to modulate the color. Pixels with no valid planes were rendered black. Voxel opacity was weighted by the scaled brightness and the fraction of valid input pixels in each bin. The volume was rendered orthographically using front-to-back compositing with a 1 µm ray step and a shared opacity multiplier of 0.04.

### Quantification of intracellular UDP-hexose by HILIC-MS

HEK293T cells grown to near-full confluency in 12-well plates were treated with 30 µM Yoda1 for 2 h or with DMSO vehicle as controls. Cells were washed three times with ice-cold DPBS (400 µL), scraped into 400 µL of cold DPBS, and lysed by brief sonication on ice. Metabolites were extracted by adding 1 mL of cold ethanol and 5 µL of internal standard (1 mM ^13^C_6_-UDP-Glc), followed by centrifugation at 12,000 × g for 20 min at 4 °C. Supernatants were collected, diluted with 4 mL of ddH_2_O, lyophilized, and reconstituted in 300 µL of acetonitrile and ddH_2_O (80:20; v:v) mixture, after which insoluble materials were removed by centrifugation at 12,000 × g for 20 min. Aliquots (30 µL) were injected into a Waters LC-MS system equipped with a SeQuant ZIC-pHILIC column (2.1 × 150 mm, 5 µm; EMD Millipore) and a Waters SQ Detector 2 single quadrupole mass spectrometer. Chromatographic separation was performed at 0.19 mL/min using mobile phase A (15 mM ammonium bicarbonate, 1 µM citric acid, pH 9 in 80% acetonitrile: 20% ddH_2_O, v:v) and mobile phase B (15 mM ammonium bicarbonate, 1 µM citric acid, pH 9 in ddH_2_O) with a 50-min gradient program (0–2 min, 100% A; 2–6 min, 100–50% A; 6–33 min, 50–20% A; 33–38 min, 20% A; 38–43 min, 20–100% A; 43–50 min, 100% A). Detection was carried out in negative electrospray ionization mode using selected-ion monitoring (SIM) at *m*/*z* 565 and 571. The endogenous signal at m/z 565 was designated UDP-hexose (UDP-Hex). For each injection, coelution of the endogenous UDP-hexose and ^13^C_6_-UDP-Glc signals defined the retention-time window used for analysis. SIM signal intensities at m/z 565 and 571 within this window were used to calculate the endogenous UDP-hexose-to-internal-standard ratio.

### Statistics and reproducibility

GraphPad Prism 10 was used for visualization and statistical analysis. Sample sizes, replicate definitions, statistical tests, and reported *P* values are given in the figure legends or figures. Unless otherwise indicated, data are presented as mean ± s.d.; mouse muscle time courses are shown as mean ± s.e.m. A value of *P* < 0.05 was considered statistically significant. No statistical method was used to predetermine sample size; sample sizes were based on prior experience with similar experiments. No blinding was performed, but samples were generally assigned to experimental groups at random. Exclusion criteria included failed transfection or viral expression, poor cell viability, imaging artifacts, or unsuccessful sensor expression. No additional data were excluded unless stated.

## Use of AI-assisted tools

The authors used ChatGPT (OpenAI), Claude (Anthropic), and Gemini (Google) to assist with literature exploration, rewording, text organization, grammar editing, and selection of statistical methods. All AI-assisted content was critically reviewed, verified, and revised by the authors, who take responsibility for the accuracy, integrity, and final content of the manuscript.

## Data availability

All data are included in the manuscript and its Supplementary Information. Source data will be provided with this paper.

## Materials availability

Key UGluS and dUGluS plasmids are being deposited to Addgene and will be available upon publication. Additional unique materials and reagents will be made available by the corresponding author upon reasonable request, subject to institutional policies and a materials transfer agreement where applicable. Requests should be directed to Hui-wang Ai.

## Acknowledgements

This work was supported by University of Virginia startup funds and NIH grants U01CA230817, R01EB033172, R01EB035430, and R01AG077773 to H.A. Photophysical characterization used resources supported by NIH BRAIN Initiative grant U24NS109107-08 to M.D.

## Author contributions

H.A. and Z.L. conceptualized the study. H.A. obtained funding, supervised the research, and provided intellectual guidance. Z.L. engineered and characterized the sensor. D.H. generated and maintained the transgenic zebrafish lines and guided Z.L. in larval imaging experiments. O.M. performed mouse tissue and live-animal imaging with assistance from Z.L. and F.H. Photophysical characterization was performed by M.D. and A.T., using protein prepared by Z.L. S.H. assisted with light-sheet zebrafish imaging. D.M.P. supervised D.H. Z.L., O.M., M.D., and S.H. analyzed the data. Z.L. and H.A. prepared the figures and wrote the manuscript, with sections provided by D.H., O.M., and M.D. and input from all authors.

## References

1. Caron, P. et al. A liquid chromatography-mass spectrometry assay for the quantification of nucleotide sugars in human plasma and urine specimens and its clinical application. J. Chromatogr. A 1677, 463296 (2022).

2. Sadoine, M., Reger, M., Wong, K.M. & Frommer, W.B. Affinity Series of Genetically Encoded Förster Resonance Energy-Transfer Sensors for Sucrose. ACS Sens. 6, 1779–1784 (2021).

3. Gest, A.M.M. et al. Molecular Spies in Action: Genetically Encoded Fluorescent Biosensors Light up Cellular Signals. Chem. Rev. 124, 12573–12660 (2024).

4. Cameron, W.D. et al. Apollo-NADP(+): a spectrally tunable family of genetically encoded sensors for NADP(+). Nat. Methods 13, 352–358 (2016).

5. Zhao, F.L., Zhang, C., Zhang, C., Tang, Y. & Ye, B.C. A genetically encoded biosensor for in vitro and in vivo detection of NADP(+). Biosens Bioelectron 77, 901–906 (2016).

6. Wei, Q. et al. A high-performance fluorescent sensor spatiotemporally reveals cell-type specific regulation of intracellular adenosine in vivo. Nat. Commun. 16, 4245 (2025).

7. Li, Z., Zhang, J. & Ai, H.-w. Genetically Encoded Green Fluorescent Biosensors for Monitoring UDP-GlcNAc in Live Cells. ACS Cent. Sci. 7, 1763–1770 (2021).

8. Mikkola, S. Nucleotide Sugars in Chemistry and Biology. Molecules 25, 5755 (2020).

9. Cifuente, J.O., Colleoni, C., Kalscheuer, R. & Guerin, M.E. Architecture, Function, Regulation, and Evolution of α-Glucans Metabolic Enzymes in Prokaryotes. Chem. Rev. 124, 4863–4934 (2024).

10. Reily, C., Stewart, T.J., Renfrow, M.B. & Novak, J. Glycosylation in health and disease. Nat. Rev. Nephrol. 15, 346–366 (2019).

11. Flynn, R.A. et al. Small RNAs are modified with N-glycans and displayed on the surface of living cells. Cell 184, 3109–3124 (2021).

12. Egger, S., Chaikuad, A., Kavanagh, K.L., Oppermann, U. & Nidetzky, B. Structure and mechanism of human UDP-glucose 6-dehydrogenase. J. Biol. Chem. 286, 23877–23887 (2011).

13. Chen, J. et al. Hepatic glycogenesis antagonizes lipogenesis by blocking S1P via UDPG. Science 383, eadi3332 (2024).

14. Kozlov, G. & Gehring, K. Calnexin cycle – structural features of the ER chaperone system. FEBS J. 287, 4322–4340 (2020).

15. Chekeni, F.B. et al. Pannexin 1 channels mediate ‘find-me’ signal release and membrane permeability during apoptosis. Nature 467, 863–867 (2010).

16. Medina, C.B. et al. Metabolites released from apoptotic cells act as tissue messengers. Nature 580, 130–135 (2020).

17. Qian, C. et al. Localization, proteomics, and metabolite profiling reveal a putative vesicular transporter for UDP-glucose. eLife 10, e65417 (2021).

18. Lazarowski, E.R. & Harden, T.K. UDP-Sugars as Extracellular Signaling Molecules: Cellular and Physiologic Consequences of P2Y_14_ Receptor Activation. Mol. Pharmacol. 88, 151–160 (2015).

19. Parandeh, F. et al. Inhibitory effect of UDP-glucose on cAMP generation and insulin secretion. J. Biol. Chem. 295, 15245–15252 (2020).

20. Flores-Diaz, M. et al. A Cellular UDP-glucose Deficiency Causes Overexpression of Glucose/Oxygen-regulated Proteins Independent of the Endoplasmic Reticulum Stress Elements J. Biol. Chem. 279, 21724–21731 (2004).

21. Doshi, M.B. et al. Disruption of sugar nucleotide clearance is a therapeutic vulnerability of cancer cells. Nature 623, 625–632 (2023).

22. Wang, X. et al. UDP-glucose accelerates SNAI1 mRNA decay and impairs lung cancer metastasis. Nature 571, 127–131 (2019).

23. Perenthaler, E. et al. Loss of UGP2 in brain leads to a severe epileptic encephalopathy, emphasizing that bi-allelic isoform-specific start-loss mutations of essential genes can cause genetic diseases. Acta Neuropathol. 139, 415–442 (2020).

24. Wolfe, A.L. et al. UDP-glucose pyrophosphorylase 2, a regulator of glycogen synthesis and glycosylation, is critical for pancreatic cancer growth. Proc. Natl. Acad. Sci. U. S. A. 118, e2103592118 (2021).

25. Zhang, X. et al. UGP2 is a potential target for breast cancer, based on bioinformatics analysis and in vitro experiments. Sci. Rep. 15, 42990 (2025).

26. Battistone, M.A., Mendelsohn, A.C., Brown, D. & Breton, S. Urinary UDP-Glucose as a Novel Actionable Biomarker of Dehydration-Induced Acute Kidney Injury. Ann. Nutr. Metab. 77, 25–27 (2021).

27. Battistone, M.A. et al. Proinflammatory P2Y14 receptor inhibition protects against ischemic acute kidney injury in mice. J. Clin. Invest. 130, 3734–3749 (2020).

28. Kim, H. et al. Structural basis for the reaction mechanism of UDP-glucose pyrophosphorylase. Mol. Cells 29, 397–405 (2010).

29. Zhang, H. et al. An Ultraresponsive Green Biosensor for Robust in Vivo Imaging of Synaptic Zinc Dynamics. ACS Sens. 11, 2398–2408 (2026).

30. Skinner, O.S. et al. Salvage of ribose from uridine or RNA supports glycolysis in nutrient-limited conditions. Nat. Metab. 5, 765–776 (2023).

31. Nwosu, Z.C. et al. Uridine-derived ribose fuels glucose-restricted pancreatic cancer. Nature 618, 151–158 (2023).

32. Bruzzone, R., Hormuzdi, S.G., Barbe, M.T., Herb, A. & Monyer, H. Pannexins, a family of gap junction proteins expressed in brain. Proc. Natl. Acad. Sci. U. S. A. 100, 13644–13649 (2003).

33. Narahari, A.K. et al. ATP and large signaling metabolites flux through caspase-activated Pannexin 1 channels. eLife 10, e64787 (2021).

34. Hansen, D.B., Braunstein, T.H., Nielsen, M.S. & MacAulay, N. Distinct permeation profiles of the connexin 30 and 43 hemichannels. FEBS Lett. 588, 1446–1457 (2014).

35. Botello-Smith, W.M. et al. A mechanism for the activation of the mechanosensitive Piezo1 channel by the small molecule Yoda1. Nat. Commun. 10, 4503 (2019).

36. Zhao, Y. et al. An expanded palette of genetically encoded Ca²⁺ indicators. Science 333, 1888–1891 (2011).

37. Cinar, E. et al. Piezo1 regulates mechanotransductive release of ATP from human RBCs. Proc. Natl. Acad. Sci. U. S. A. 112, 11783–11788 (2015).

38. Zeng, Y. et al. Mechanosensitive piezo1 calcium channel activates connexin 43 hemichannels through PI3K signaling pathway in bone. Cell Biosci. 12, 191 (2022).

39. Kwan, K.M. et al. The Tol2kit: A multisite gateway-based construction kit for Tol2 transposon transgenesis constructs. Dev. Dyn. 236, 3088–3099 (2007).

40. Dane, P.J. & Tucker, J.B. Modulation of epidermal cell shaping and extracellular matrix during caudal fin morphogenesis in the zebra fish Brachydanio rerio. J. Embryol. Exp. Morphol. 87, 145–161 (1985).

41. Lalonde, R.L. & Akimenko, M.A. Effects of fin fold mesenchyme ablation on fin development in zebrafish. PLoS One 13, e0192500 (2018).

42. Zimmer, B.M., Barycki, J.J. & Simpson, M.A. Integration of Sugar Metabolism and Proteoglycan Synthesis by UDP-glucose Dehydrogenase. J. Histochem. Cytochem. 69, 13–23 (2021).

43. Yan, Z., Yan, H. & Ou, H. Human thyroxine binding globulin (TBG) promoter directs efficient and sustaining transgene expression in liver-specific pattern. Gene 506, 289–294 (2012).

44. Fleig, W.E., Enderle, D., Steudter, S., Nöther-Fleig, G. & Ditschuneit, H. Regulation of basal and insulin-stimulated glycogen synthesis in cultured hepatocytes. Inverse relationship to glycogen content. J. Biol. Chem. 262, 1155–1160 (1987).

45. Nozaki, Y. et al. Metabolic control analysis of hepatic glycogen synthesis in vivo. Proc. Natl. Acad. Sci. U. S. A. 117, 8166–8176 (2020).

46. Salva, M.Z. et al. Design of tissue-specific regulatory cassettes for high-level rAAV-mediated expression in skeletal and cardiac muscle. Mol. Ther. 15, 320–329 (2007).

47. Sylow, L., Tokarz, V.L., Richter, E.A. & Klip, A. The many actions of insulin in skeletal muscle, the paramount tissue determining glycemia. Cell Metab. 33, 758–780 (2021).

48. Jensen, J., Rustad, P.I., Kolnes, A.J. & Lai, Y.C. The role of skeletal muscle glycogen breakdown for regulation of insulin sensitivity by exercise. Front. Physiol. 2, 112 (2011).

49. Adeva-Andany, M.M., González-Lucán, M., Donapetry-García, C., Fernández-Fernández, C. & Ameneiros-Rodríguez, E. Glycogen metabolism in humans. BBA Clin. 5, 85–100 (2016).

50. Parichy, D.M., Ransom, D.G., Paw, B., Zon, L.I. & Johnson, S.L. An orthologue of the kit-related gene fms is required for development of neural crest-derived xanthophores and a subpopulation of adult melanocytes in the zebrafish, Danio rerio. Development 127, 3031–3044 (2000).

51. Moffat, J. et al. A lentiviral RNAi library for human and mouse genes applied to an arrayed viral high-content screen. Cell 124, 1283–1298 (2006).

52. Drobizhev, M., Makarov, N.S., Tillo, S.E., Hughes, T.E. & Rebane, A. Two-photon absorption properties of fluorescent proteins. Nat. Methods 8, 393–399 (2011).

53. Barnett, L.M., Hughes, T.E. & Drobizhev, M. Deciphering the molecular mechanism responsible for GCaMP6m’s Ca^2+^-dependent change in fluorescence. PLoS One 12, e0170934 (2017).

54. Drobizhev, M., Molina, R.S. & Hughes, T.E. Characterizing the Two-photon Absorption Properties of Fluorescent Molecules in the 680-1300 nm Spectral Range. Bio-Protoc. 10, e3498 (2020).

55. Suster, M.L., Kikuta, H., Urasaki, A., Asakawa, K. & Kawakami, K. Transgenesis in zebrafish with the tol2 transposon system. Methods Mol. Biol. 561, 41–63 (2009).

56. Schindelin, J. et al. Fiji: an open-source platform for biological-image analysis. Nat. Methods 9, 676–682 (2012).

57. Kardash, E., Bandemer, J. & Raz, E. Imaging protein activity in live embryos using fluorescence resonance energy transfer biosensors. Nat. Protoc. 6, 1835–1846 (2011).

