## Supplement Information for "Repurposing a biosynthetic enzyme for imaging UDP-glucose dynamics in living systems"

**Supplementary Information**

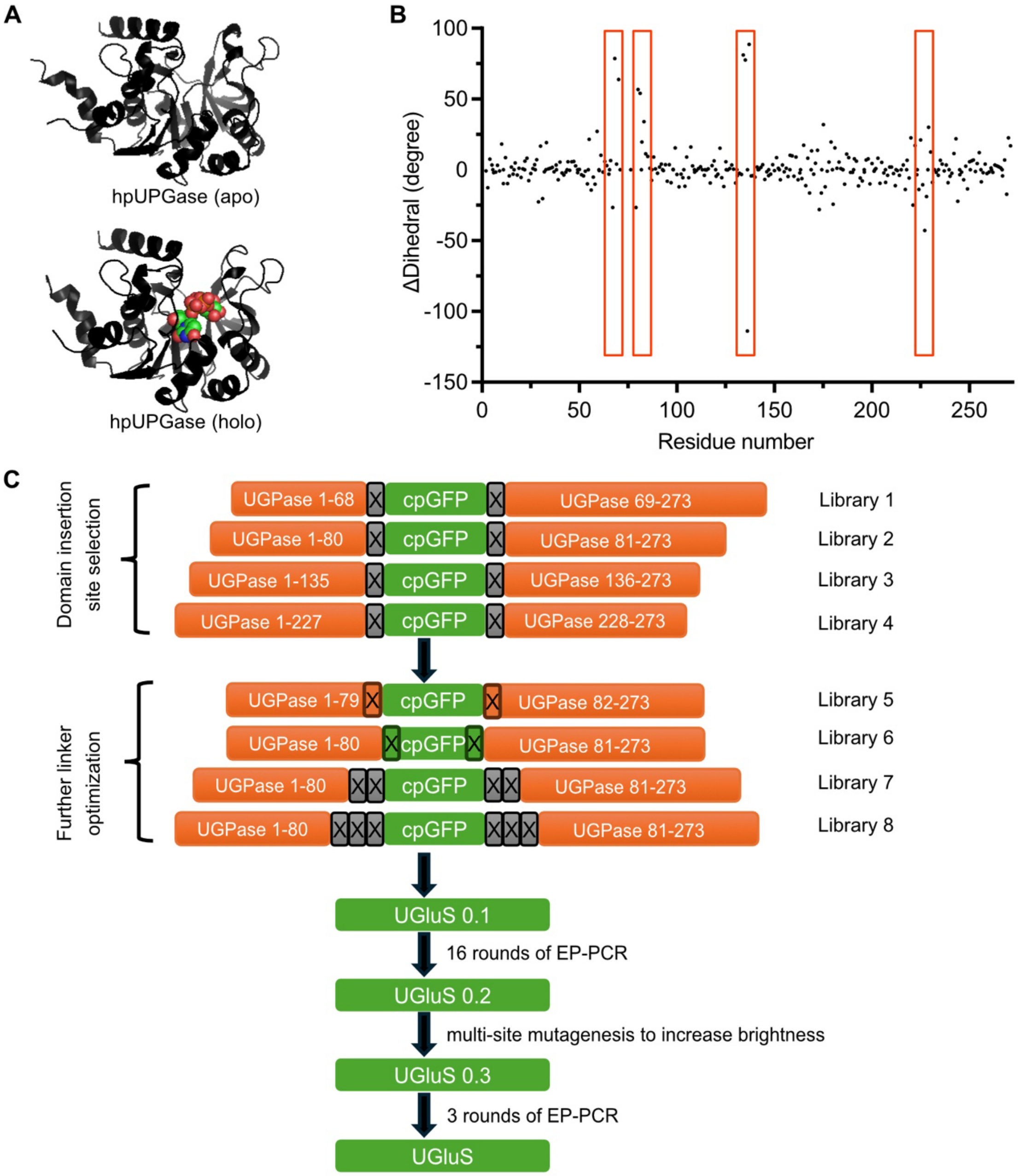


**Supplementary Fig. 1. Structural comparison and engineering workflow for UGluS.**

(**A**) Crystal structures of *Helicobacter pylori* UDP-glucose pyrophosphorylase (HpUGPase) in the apo state (PDB 3JUJ) and UDP-Glc/$Mg^{2+}$-bound state (PDB 3JUK).

(**B**) Differences in Cα pseudo-dihedral angles, defined by four consecutive Cα atoms, between the UDP-Glc-bound and apo structures. Orange boxes mark candidate cpGFP insertion regions.

(**C**) UGluS engineering workflow comprising cpGFP insertion-site screening, linker optimization, 16 rounds of error-prone PCR (EP-PCR), mutagenesis to increase fluorescence brightness, and three additional EP-PCR rounds. X denotes a randomized amino acid at a sensor junction. cpGFP, circularly permuted GFP.

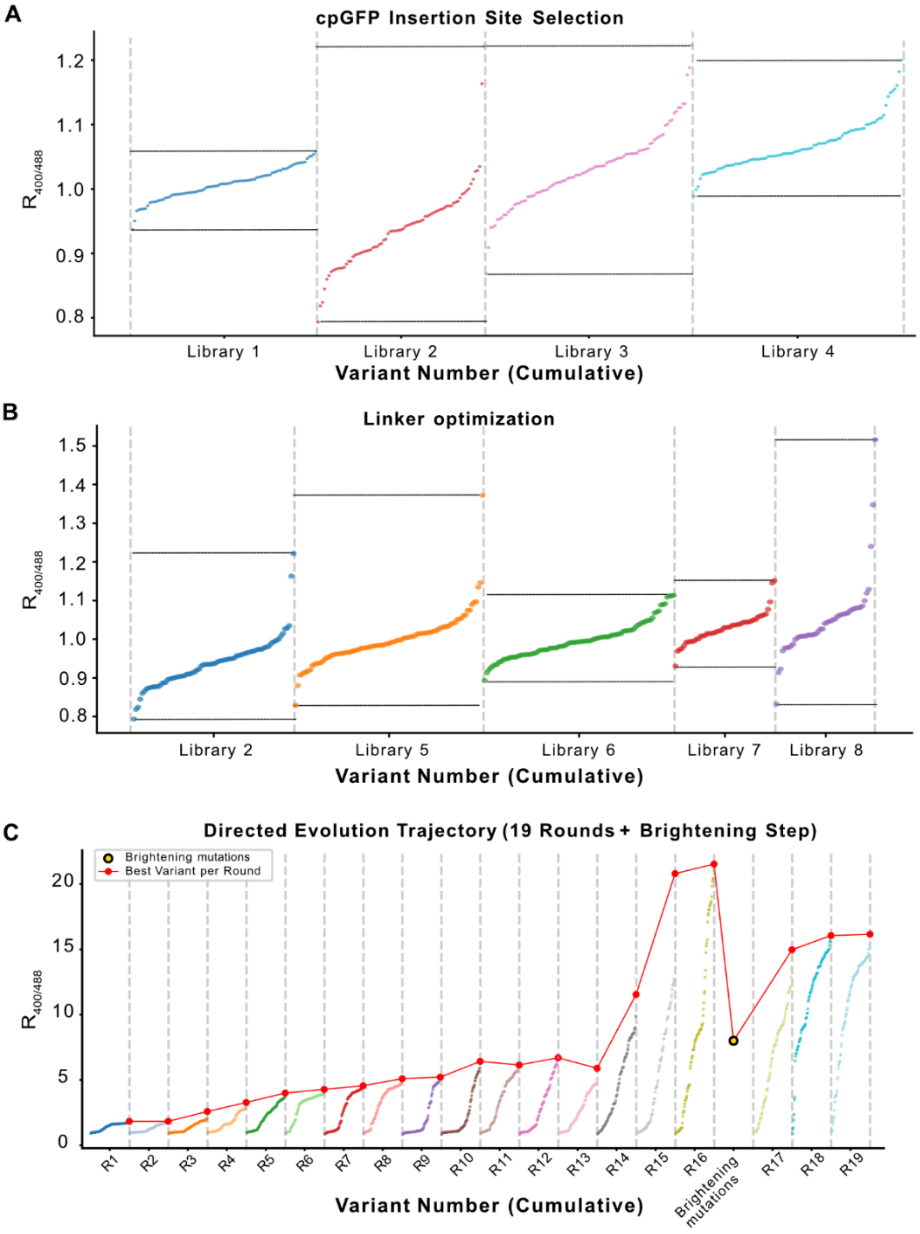


**Supplementary Fig. 2. Library screening and directed evolution of UGluS.**

(**A**) Ratiometric screening of cpGFP insertion-site libraries 1–4. Library designs are shown in Supplementary Fig. 1C.

(**B**) Ratiometric screening of linker-optimization libraries 2 and 5–8. Library 2 also appears in panel A, and the same measurements are reused in panel B.

(**C**) Screening trajectory over 19 rounds of error-prone PCR (EP-PCR), with an additional mutagenesis step to increase fluorescence brightness between rounds 16 and 17. Red markers and connecting lines identify the best variant from each round; the yellow marker identifies the variant after the brightening step.

In **A** and **B**, black horizontal lines indicate the maximum and minimum normalized fluorescence excitation ratios (400/488 nm) observed within each screened library. In **A–C**, colored points represent individual variants, ranked by their normalized fluorescence excitation ratios within each library or screening round. Each point represents a single measurement from an independently screened colony, without technical replication.

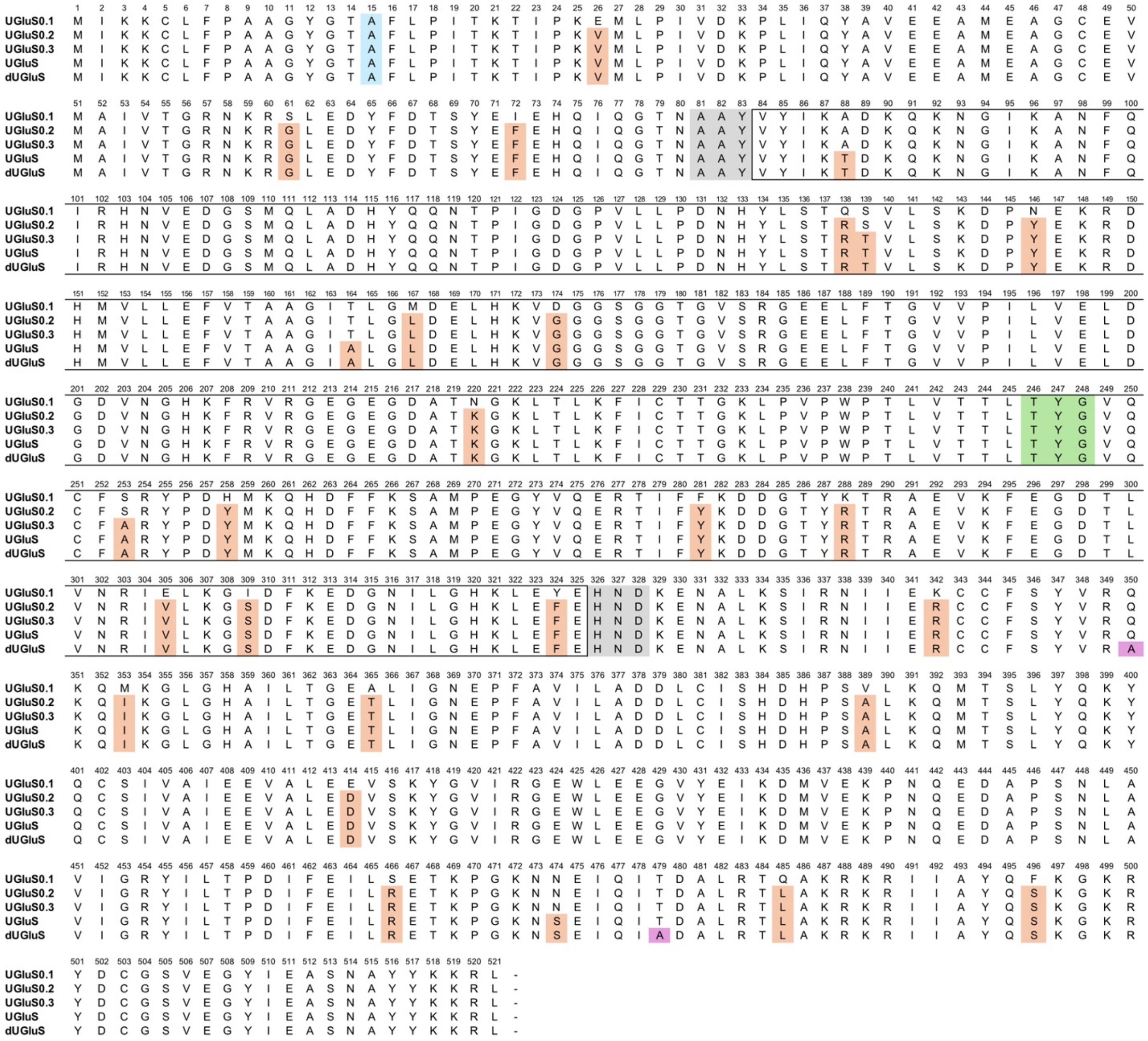


**Supplementary Fig. 3. Sequence alignment of UGluS variants and the dUGluS control.**

Amino acid sequence alignment of UGluS0.1, UGluS0.2, UGluS0.3, UGluS and dUGluS. Cyan shading marks the catalytic-inactivation site shared by all variants. Orange shading marks substitutions acquired during optimization relative to UGluS0.1; gray shading marks the junction linkers; green shading marks the chromophore-forming residues; and purple shading marks the Q350A and T479A substitutions introduced into UGluS to generate dUGluS. Black outlines delimit the cpGFP segment.

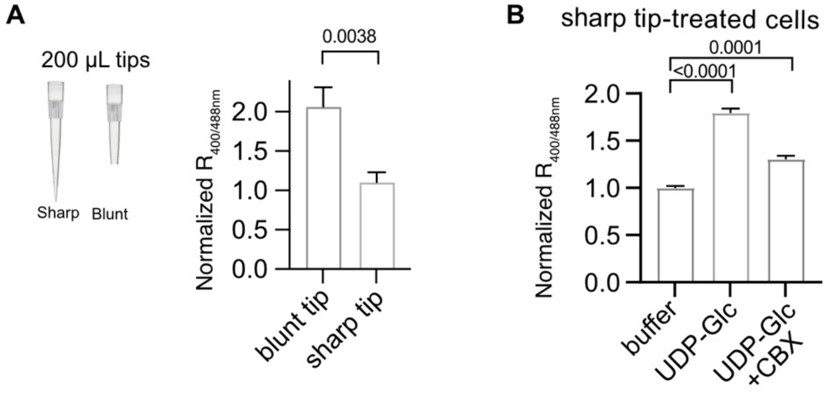


**Supplementary Fig. 4. Effects of mechanical perturbation and extracellular UDP-glucose on UGluS responses in HEK293T cells.**

(**A**) Left, illustration of sharp and blunt 200-µL pipette tips used to induce different degrees of mechanical perturbation during cell handling. Right, normalized UGluS fluorescence excitation ratios (400/488 nm) in HEK293T cells subjected to handling with sharp or blunt pipette tips.

(**B**) Normalized UGluS fluorescence excitation ratios (400/488 nm) in HEK293T cells subjected to handling with sharp pipette tips, followed by treatment with buffer alone, extracellular UDP-glucose (UDP-Glc), or UDP-Glc in the presence of carbenoxolone (CBX).

Data are mean ± SD ($n$ = 3 replicate wells per group). $P$ values were determined using an unpaired, two-tailed Student's $t$-test (**A**) or one-way ANOVA with Dunnett's multiple-comparisons test (**B**).

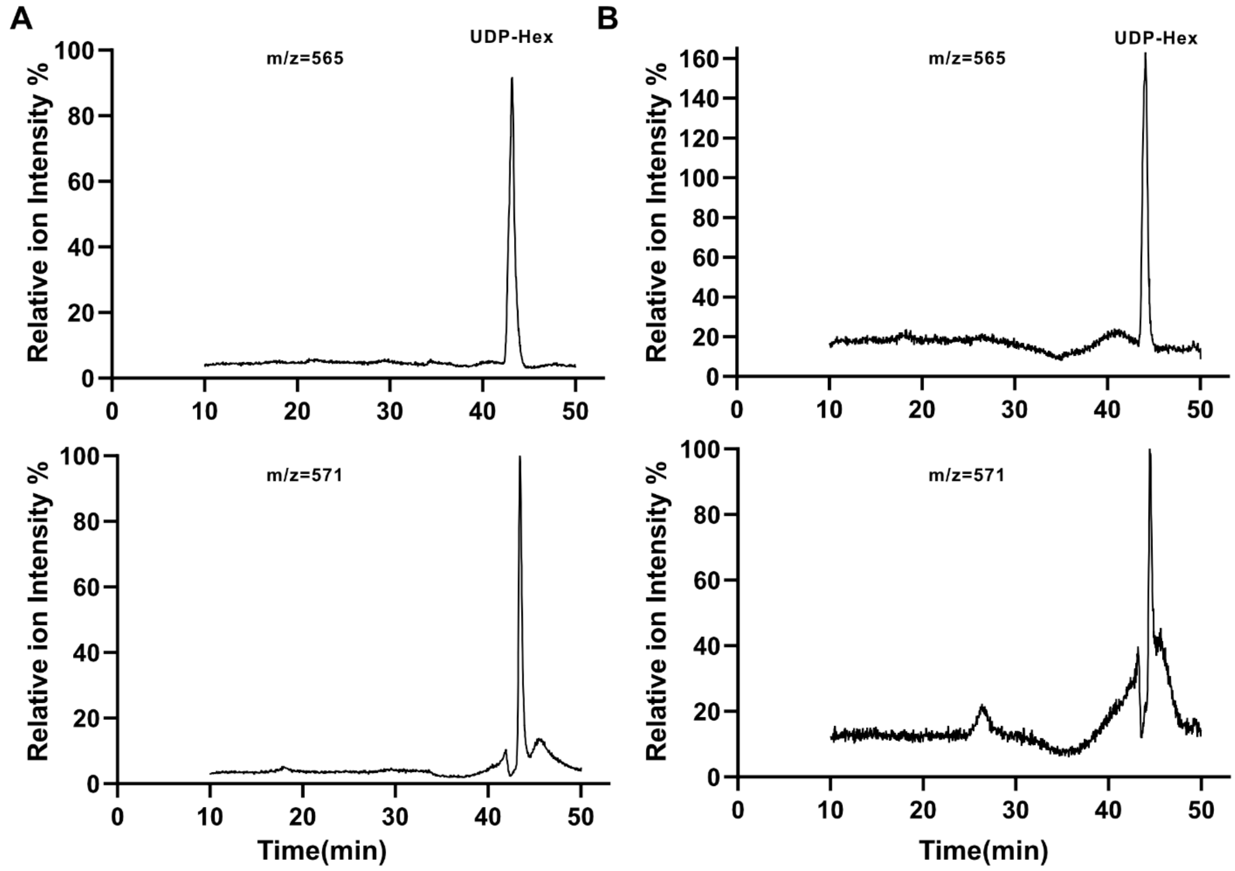


**Supplementary Fig. 5. HILIC-MS analysis of intracellular UDP-hexose in HEK293T cells.**

(**A, B**) Representative selected-ion monitoring (SIM) chromatograms of extracts from DMSO-treated control (**A**) and Yoda1-treated (**B**) HEK293T cells, showing signals at *m/z* 565 (endogenous UDP-hexose) and *m/z* 571 ($^{13}C_6$-UDP-glucose internal standard). Ion intensities were normalized to the maximum intensity of the internal-standard signal within each injection, defined as 100%. The endogenous UDP-hexose signal coeluted with the isotope-labeled internal standard within each injection, although absolute retention times varied between injections. The coeluting peaks were used to define a retention-time window, within which SIM signal intensities at *m/z* 565 and 571 were measured to calculate the endogenous UDP-hexose-to-internal-standard ratios used for quantification in Fig. 3D.

**Supplementary Table 1. Photophysical and UDP-Glc-response properties of UGluS.**

| | **UGluS** | |
|---|---|---|
| | **Ligand-free** | **Ligand-bound** |
| Excitation maxima (nm) | 398 & 488 | 398 & 493 |
| Emission maximum (nm) | 508 | 512 |
| Ratiometric response ($R/R_0$)[a] | ~18 | |
| Apparent $K_d$ (μM)[b] | 32.5 ± 1.4 | |
| Fluorescence lifetime components at 445-nm excitation (ns) | 2.61 (81%)<br>0.75 (19%) | 2.81 (72%)<br>0.66 (28%) |
| Molar extinction coefficient ($mM^{-1}$ $cm^{-1}$) | 17.3 (398 nm)<br>27.4 (490 nm) | 30.9 (398 nm)<br>5.0 (490 nm) |
| Fluorescence quantum yield | 0.59 (398 nm)<br>0.71 (490 nm) | 0.74 (398 nm)<br>0.59 (490 nm) |
| Two-photon brightness at 940 nm, $F_2$ (GM) | 7.4 | 1.84 |

[a] $R = F_{400}/F_{488}$, where $F_{400}$ and $F_{488}$ are fluorescence intensities measured with excitation at 400 and 488 nm, respectively. $R_0$ denotes the corresponding ligand-free ratio.

[b] The apparent $K_d$ was obtained from the UDP-Glc concentration–response curve (protein-based assays).

**Supplementary Table 2. Target sequences used for shRNA-mediated gene knockdown.**

| Gene/control | shRNA target sequence (5′→3′) | Reference |
|---|---|---|
| UGP2 | CCACAGCATCATCACATGAAT | https://doi.org/10.1073/pnas.2103592118 |
| PGM1 | TCGGTAGAAGCTTATGCTACA | https://doi.org/10.1371/journal.pbio.2006483 |
| Non-targeting control[a] | TTCTCCGAACGTGTCACGTAC | https://doi.org/10.1038/ncomms11848 |

[a] Negative control. The shRNA containing this sequence is referred to as shNC.

**Supplementary Table 3. Sequences of oligonucleotides used in this study.**
Oligonucleotides used for cloning, mutagenesis and gene knockdown. Sequences are written 5′ to 3′. Degenerate bases follow IUPAC notation: N, A/C/G/T; K, G/T; M, A/C; V, A/C/G; B, C/G/T.

| Oligonucleotide | Sequence (5′→3′) |
|---|---|
| galU(pbad)-F | GAGGAATTAACCATGGGCTCGAGTATGATAAAGAAATGCCTTTTTCC |
| His6(pbd)-R | ATCCGCCAAAACAGCCAAGCTTAATGATGGTGGTGATGGTG |
| galU(His6)-R | TGATGGTGGTGATGGTGGAATTCAAGTCTTTTTTTATAATAGGCATT |
| cpGFP(galU 69)-R | TGATGCTCAATTTCATAGCTMNNCTCGTACTCCAGCTTGTGC |
| galU 69-F | AGCTATGAAATTGAGCATCA |
| cpGFP(galU 68)-F | TGGAAGACTATTTCGACACANNKGTTTATATCAAGGCCGACAA |
| galU 68-R | TGTGTCGAAATAGTCTTCCAG |
| galU 81-F | AAAGAAAACGCACTCAAAAG |
| cpGFP(galU 81)-R | CTTTTGAGTGCGTTTTCTTTMNNCTCGTACTCCAGCTTGTGC |
| cpGFP(galU 80)-F | ATCAAATTCAGGGAACCAATNNKGTTTATATCAAGGCCGACAA |
| galU 80-R | ATTGGTTCCCTGAATTTGAT |
| cpGFP(galU 136)-R | AGCACAGAAGGGTGATCGTGMNNCTCGTACTCCAGCTTGTG |
| galU 136-F | CACGATCACCCTTCTGTG |
| cpGFP(galU 135)-F | CCGATGATCTGTGTATTTCANNKGTTTATATCAAGGCCGACAA |
| galU 135-R | TGAAATACACAGATCATCGGC |
| cpGFP(galU 228)-R | TCTGTAATCTGAATCTCGTTMNNCTCGTACTCCAGCTTGTG |
| galU 228-F | AACGAGATTCAGATTACAGAC |
| cpGFP(galu 227)-F | AGACTAAGCCAGGCAAGAATNNKGTTTATATCAAGGCCGACAA |
| galU 227-R | ATTCTTGCCTGGCTTAGTC |
| cpGFP 3VVC(gal80)-R | CTTTTGAGTGCGTTTTCTTTGBBGBBGBBCTCGTACTCCAGCTTGTGCC |
| cpGFP 2VVC(gal80)-R | CTTTTGAGTGCGTTTTCTTTGBBGBBCTCGTACTCCAGCTTGTGC |
| cpGFP 3VVC (gaul80)-F | ATCAAATTCAGGGAACCAATVVCVVCVVCGTTTATATCAAGGCCGACAA |
| cpGFP 2VVC (gaul80)-F | ATCAAATTCAGGGAACCAATVVCVVCGTTTATATCAAGGCCGACAA |
| cpGFP(galU82 NNK)-R | ATGCTTTTGAGTGCGTTTTCMNNCTCGTACTCCAGCTTGTGC |
| cpGFP NNK(gal81)-R | CTTTTGAGTGCGTTTTCTTTMNNGTACTCCAGCTTGTGCCC |
| cpGFP(galu79 NNK)-F | AGCATCAAATTCAGGGAACCNNKGTTTATATCAAGGCCGACAA |
| cpGFP NNK (galu80)-F | ATCAAATTCAGGGAACCAATNNKTATATCAAGGCCGACAAGC |
| galU 82-F | GAAAACGCACTCAAAAGCA |
| galU 79-R | GGTTCCCTGAATTTGATGC |

| Oligonucleotide | Sequence (5′→3′) |
| --- | --- |
| galU(pcDNA3)-F | CTCACTATAGGGAGACCCAAGCTTGCCACCATGATAAAGAAATG CCTTTTTCC |
| His6(pcd3)-F | CACTATAGGGAGACCCAAGCTTGCCACCCATCATCATCAT |
| His6(plenti)-R | GAATTCCTGCAGCCCGTAGTTCTAGATTAATGATGGTGGTGATG GT |
| galU(plent)-F | GACTAGCCTCGAGGTTTGCTAGCGGCACCATGATAAAGAAATG |
| galU(mhck7)-F | GTCCAGGCGGCCGCTCTAGAGCCACCATGATAAAGAAATG |
| His6(mhck7)-R | GTTGATTGGATCCGGGTACCTTAATGATGGTGGTGATGGT |
| His6(TBG)-R | AGGTTGATTGGATCCAAGCTTAATGATGGTGGTGATGGT |
| galU(TBG)-F | ACAGGTGTCCAGGCGGCCGCGCCACCATGATAAAGAAATG |
| pLKO.1-shNC-F | CCGGTTCTCCGAACGTGTCACGTACCTCGAGGTACGTGACACGT TCGGAGAATTTTTG |
| pLKO.1-shNC-R | AATTCAAAAATTCTCCGAACGTGTCACGTACCTCGAGGTACGTG ACACGTTCGGAGAA |
| shUGP2-F | CCGG CCACAGCATCATCACATGAAT CTCGAG ATTCATGTGATGATGCTGTGG TTTTTG |
| shUGP2-R | AATTCAAAAA CCACAGCATCATCACATGAAT CTCGAG ATTCATGTGATGATGCTGTGG |
| shPGM1-F | CCGG TCGGTAGAAGCTTATGCTACA CTCGAG TGTAGCATAAGCTTCTACCGA TTTTTG |
| shPGM1-R | AATTCAAAAA TCGGTAGAAGCTTATGCTACA CTCGAG TGTAGCATAAGCTTCTACCGA |
| UG-Q102A-R | CAGCCCTTTTATCTGCTTAGCTCTGACATAACTAAAGCA |
| UG-103K-F | AAGCAGATAAAAGGGCTG |
| UG-T231A-R | CAGTGTCCGCAGAGCGTCAGCAATCTGAATCTCGTTATTC |
| UG-D232-F | GACGCTCTGCGGACACTG |

**Supplementary Video 1 | Ratiometric UGluS response to glucose supplementation in HEK293T cells.**

Time-lapse fluorescence imaging of UGluS in HEK293T cells following stimulation with 25 mM glucose. Scale bar, 20 μm.

**Supplementary Video 2 | Ratiometric UGluS response to uridine supplementation in HEK293T cells.**

Time-lapse fluorescence imaging of UGluS in HEK293T cells following stimulation with 20 mM uridine. Scale bar, 20 μm.

**Supplementary Video 3 | Concurrent UGluS and R-GECO imaging during Yoda1 stimulation in HEK293T cells.**

Time-lapse imaging of UGluS and the $Ca^{2+}$ indicator R-GECO in HEK293T cells following stimulation with 30 μM Yoda1. Scale bar, 20 μm.

**Supplementary Video 4 | UGluS response to uridine stimulation in larval zebrafish.**

Time-lapse fluorescence imaging of UGluS in zebrafish larvae following stimulation with 50 mM uridine. Scale bar, 100 μm.

**Supplementary Video 5 | Light-sheet ratiometric imaging of UGluS in larval zebrafish.**

Volumetric light-sheet ratiometric video of UGluS-expressing zebrafish larvae. Scale bar, 500 μm.
